# The Oral Microbiome Is a Population-Scale Readout of the Exposome, Age, and Systemic Health

**DOI:** 10.64898/2026.02.23.707541

**Authors:** Byeongyeon Cho, Aleksandar D. Kostic, Braden T. Tierney, Chirag J. Patel

**Affiliations:** Department of Biomedical Informatics, Harvard Medical School, Boston, MA, USA; Department of Microbiology and Immunobiology, Harvard Medical School, Boston, MA, USA; Section on Pathophysiology and Molecular Pharmacology, Joslin Diabetes Center, Boston, MA, USA; The Two Frontiers Project, Fort Collins, CO, USA

**Keywords:** microbiome, microbiota, oral microbiome, disease associations, machine learning, diet, nutrition, dysbiosis, community composition, exposome

## Abstract

The oral microbiome interfaces humans and the environment and is implicated in diseases from caries to cardiovascular conditions. Yet, few studies systematically interrogate oral taxa associations with the host phenome and exposome in diverse populations. We developed a comprehensive oral microbiome atlas, deploying a Microbiome Association Study (MAS) evaluating relationships between host features including exposome, disease, and physiology and the microbiome in a 10,000-person representative US population. Evaluating demographics, 133 phenotypes, 473 exposures, and 20 disease outcomes across 1,349 taxa yielded >800k relationships and 45,757 FDR-significant associations. Age emerged as a major organizing axis, with genera following non-linear life-course patterns. Oral disease, smoking, and dietary sugar correlated with aciduric and anaerobic taxa, whereas oral health featured oxygen-tolerant *Proteobacteria*. The exposome and cardiovascular/respiratory disease linked to diverse taxa. These results establish the oral microbiome as a sensitive, population-scale indicator of the exposome, phenome, and systemic health.

## Introduction

The human oral cavity contains a complex and diverse microbial ecosystem. These organisms span over 600 genera, comprise a range of physiologies (e.g., anaerobes and aerobes), and are arrayed in complex, spatially-delineated structures.^1–3^ With the mouth serving as a key interface between humans and environmental exposures including dietary factors, toxicants and behaviors such as smoking^4^, the composition of the oral microbiota is related to both host-intrinsic and extrinsic factors. For example, the oral microbiome is known to be associated with host phenotypes local to the mouth, such as tooth decay and gingivitis, but also hypothesized to influence systemic phenotypes, such as cardiovascular disease and inflammatory bowel disease.^5–7^ Additionally, particular species that typically colonize the oral environment, when abundant in other areas of the GI tract, are thought to be indicative of certain disease characteristics such as that of colorectal cancer.^8–10^

However, despite the importance of this system, how the oral microbiota associates with host phenotype and environmental factors across diverse populations has been understudied.^11,12^ Furthermore, there has yet to be a systematic analysis of how the oral microbiome varies along the joint axes of demographics, exposure, and host physiology. Doing so requires comprehensive cohort data that captures both the exposome (i.e., the integration of physical, psychosocial, and biological factors to which humans are exposed) and the phenome (i.e., factors that are connected to human disease and health).^13,14^ Investigations thus far on the human oral microbiome tends to focus on a small subset of factors at a time, which can limit our understanding of how the oral microbiome contributes to disease.

Unprecedented insight into population-level oral microbial dynamics is now possible through a 10,000-person oral microbiome cohort sequenced^15^ in participants of the US Centers for Disease Control (CDC) National Health and Nutrition Examination Survey (NHANES)^16^. In addition to demographic information that is representative of the entire non-institutionalized population of the United States, NHANES offers a comprehensive view of the host exposome through objective and CDC-prioritized blood (e.g, lead and cadmium), urinary (plastics and phenols), and dietary measurements (self-reported and objective indicators of nutrition). NHANES also contains clinical phenotypes that include glucose and body mass index. In sum, these provide an unparalleled database for studying human host oral microbiome relationships along both the exposomic and phenomic axes.

Previous work using this cohort has assessed demographic shifts in oral microbiome profiles and linked diversity or selected taxa abundance to rheumatoid arthritis, depression, type 2 diabetes, frailty, diet quality, and medication burden^17–23^. This body of work has primarily examined a narrow disease domain or exposure class at a time and have focused primarily on α-/β-diversity or a small number of genera, without leveraging the full breadth of NHANES phenotypic and exposomic measurements or community-level signatures, leaving the multidimensional link between host traits and diseases, environmental exposures (diet and the chemical/physical environment), and the oral microbiome largely unexplored. As a result, we still lack a systematic understanding of how host traits, behaviors, and exposures are reflected in oral microbial community structure and, crucially, how these community-level signatures relate to health and disease.

To address this gap, we conducted a systematic analysis of the oral microbiome by testing its associations with more than 600 host variables spanning demographics, clinical traits, behaviors, environmental exposures, and disease outcomes. We also introduce an ecological and functional layer of interpretation of these relationships by integrating microbiome phenotype data from the JGI GOLD database. Furthermore, we show that oral signatures associated with host trait variation are broadly distributed across a complex community of genera, with few conditions dominated by any single taxonomic group, implying that oral microbial influences on human health emerge from the concerted actions of many weak, distributed associations rather than from a few dominant microbial determinants.

## Results

### Host demographic traits are correlated with distinct microbial community structures

To characterize the composition of the U.S. oral microbiome, we quantified the most abundant taxa across taxonomic levels from phylum to genus (Figure 1A; Supplementary Figure 1). The U.S. oral microbiome is dominated by *Firmicutes*, *Bacteroidetes*, *Actinobacteria*, *Proteobacteria*, and *Fusobacteria*, with 98% of individuals harboring microbes from these phyla. At the genus level, the predominance of *Firmicutes* (51% relative abundance) was driven largely by *Streptococcus*, which accounted for 32.3% of total abundance. Additionally, in the NHANES cohort, we observed marked divergence between the reported prevalence^17^ (non-zero detection) and relative abundance; taxa that are widely prevalent can be present at low abundance, such as members of the *Saccharibacteria* phylum.

**Figure 1.**
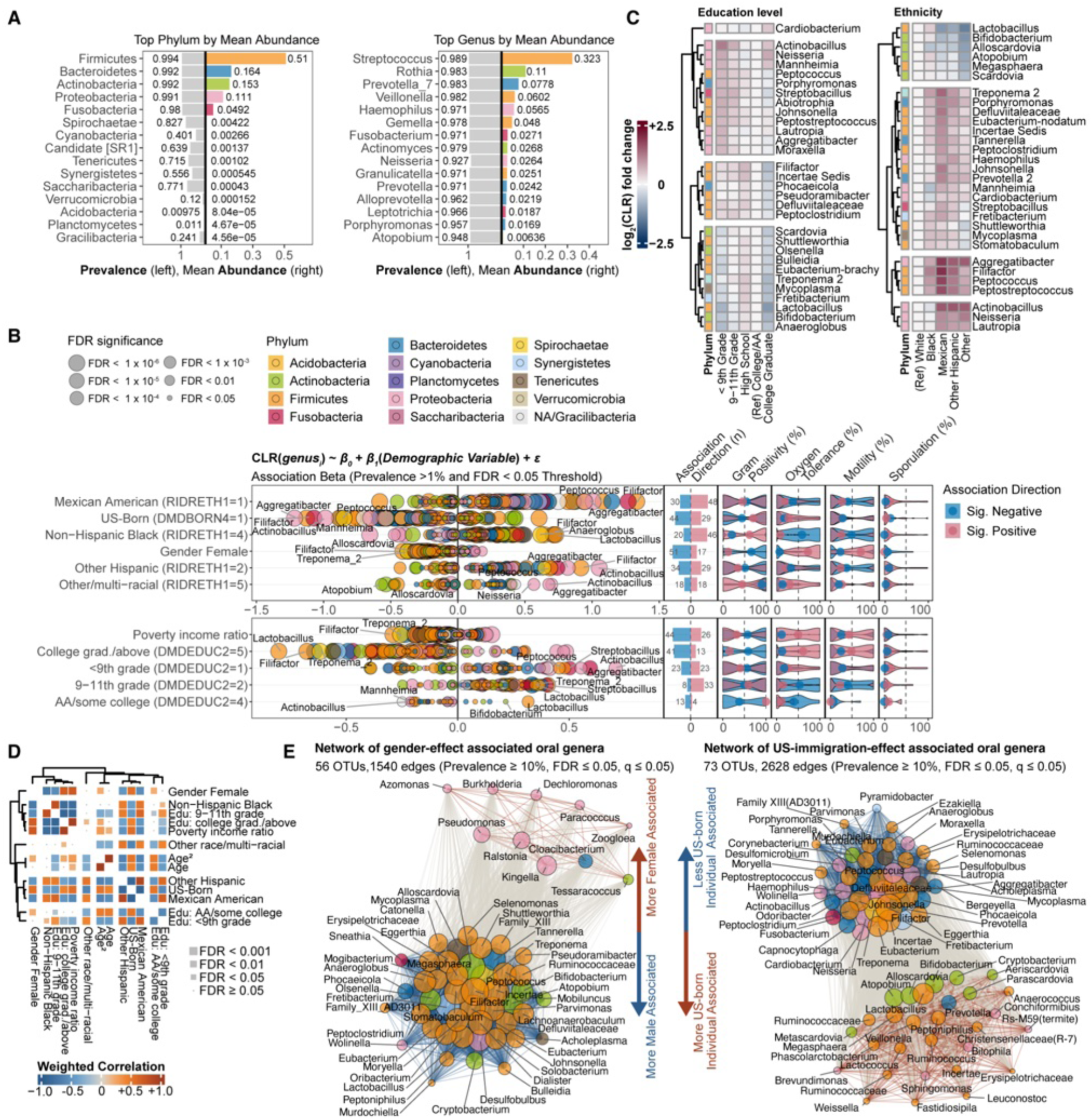
Composition and demographic associations of the US oral microbiome. (A) Phylum- and genus-level composition of the oral microbiome. The biplot displays the mean relative abundance of the top 15 most abundant phyla (left) and genera (right), alongside their prevalence across participants. Phylum-level color coding is consistent across all plots. (B) Survey-weighted associations of 1,349 CLR-transformed oral genera with 11 demographic variables across two NHANES cycles (2009–2010, 2011–2012). Left: x-axis shows regression coefficients (restricted to OTUs with prevalence ≥ 1%, FDR ≤ 0.05 and q ≤ 0.05), with dot size indicating FDR significance. Right: GOLD-DB–derived microbial phenotype indices (gram positivity, oxygen tolerance, motility, sporulation; see Methods) summarized by association direction (positive/negative) and restricted to taxa with complete annotation. (C) The top 30 most significantly variable (Kruskal–Wallis FDR ≤ 0.05) genera (≥1% prevalence) by CLR log₂ fold change (capped at –2.5 to +2.5); dendrograms show Euclidean distance. (D) Microbial association signatures across 13 demographic variables. Each cell represents the inverse-variance–weighted Pearson correlation between pairs of survey-weighted OTU-level effect (β) estimates derived from CLR-transformed svyglm models, restricted to variables with at least two significantly associated (FDR < 0.05) common microbes (prevalence > 10%). Cell size indicates FDR significance (≤ 0.001, ≤ 0.01, ≤ 0.05, > 0.05). (E) Small-effect network visualization of gender (left) and US-born (right) effects from common (prevalence ≥ 10%) community of oral genera (FDR ≤ 0.05, q ≤ 0.05). Node size scales with cumulative effect magnitude; color denotes phylum. Edges link genera with concordant associations (similar-effect microbes closer; dissimilar microbes further); width reflects geometric mean of effect sizes; color encodes direction (blue = negative, orange = positive) (see Methods).

Building on previous studies’ ^18^ findings of sex, race, and socioeconomic associations to genus-level composition, we statistically evaluated the demographic associations with oral microbiome composition by testing 13 selected variables against 1,349 genera (17,537 tests total; Table 1) using survey-weighted approaches to account for the sampling of the NHANES. After confirming these demographic factors as important covariates (Fig 1,2), we constructed and ran four regression schemas (see Methods; Table 1) to associate 626 host variables with centered log-ratio (CLR)–transformed genus abundances (Supplementary Table S3 reports results for non-normalized abundances). Supplementary Table 1 provides the full set of variable definitions and their distributions across NHANES participants, and complete association results and summary statistics are reported in Supplementary Table S2. All of the following associations reported in this and other sections were significant after adjusting for multiple hypothesis testing at the false discovery rate (FDR) 5% correction level via Benjamini-Hochberg adjustment. We describe the percentage and total number of results for different FDR approaches in Table 1.

**Figure 2.**
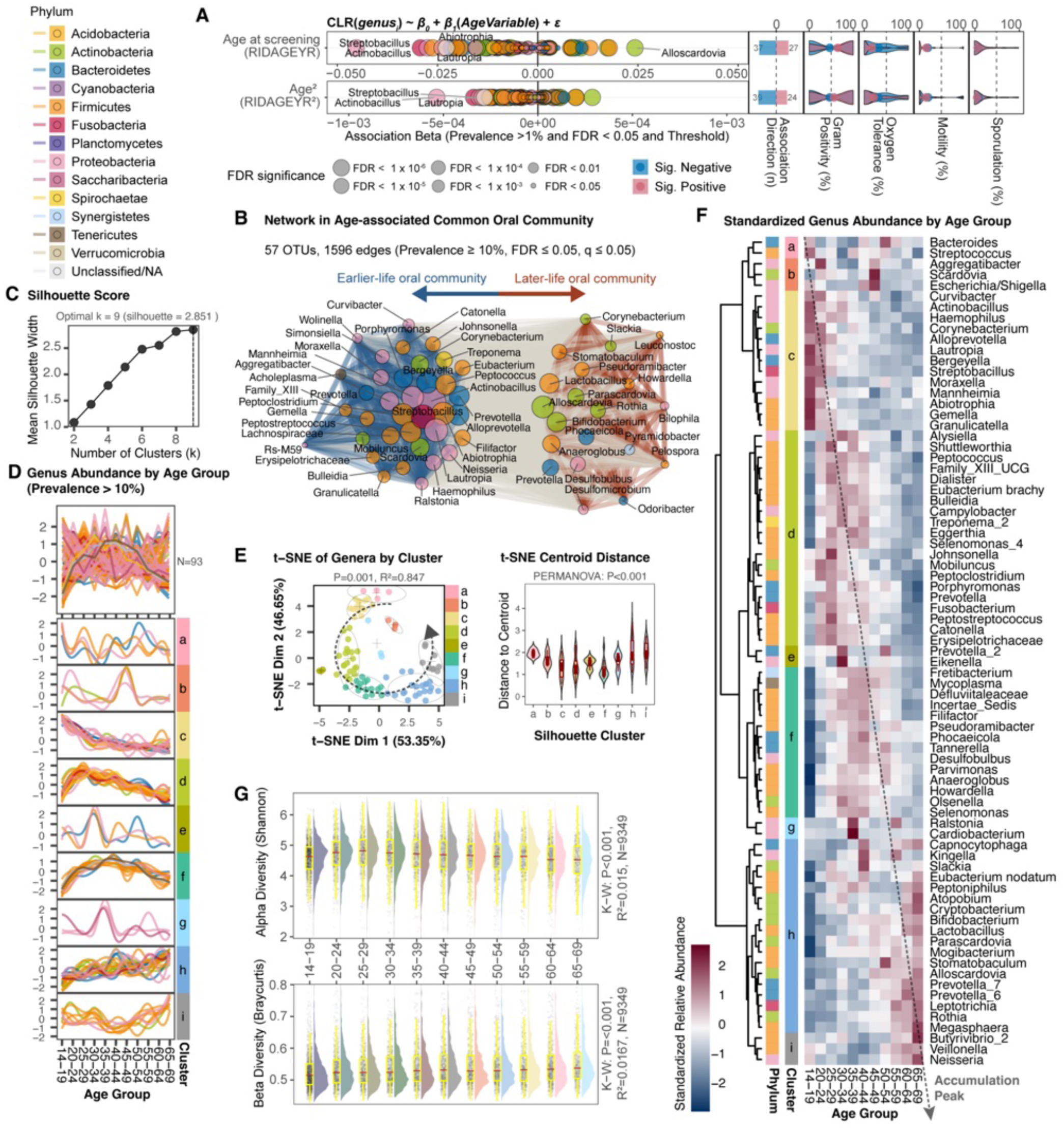
Age-driven nonlinear accumulation patterns in the oral microbiome trace a pseudotime-like community gradient. (A) Survey-weighted associations of 1,349 CLR-transformed OTUs on age and age² across two NHANES cycles (2009–2010, 2011–2012). Left: regression coefficients. Right: GOLD-DB–derived microbial phenotype indices (oxygen tolerance, motility, sporulation) by significant association direction (positive/negative) restricted to taxa with complete annotation. (B) Network of age-associated common (prevalence ≥ 10%) community of oral genera (FDR ≤ 0.05, q ≤ 0.05). Node size scales with cumulative effect magnitude; color denotes phylum. Edges link genera with concordant age associations; width reflects geometric mean of effect sizes; color encodes direction (blue = negative, orange = positive). (C) Cluster determination diagnostics showing mean silhouette width versus number of clusters (k = 2–9); dashed line marks optimal k = 9. (D) Age-trajectory clustering of 93 prevalent genera (≥10% of participants) across 11 age groups (14–19 to 65–69 years). Top: standardized (z-score) mean abundances; bottom: nine silhouette-defined clusters (k = 9; silhouette = 2.851) with trajectories colored by phylum (cluster sizes: a = 4, b = 4, c = 14, d = 20, e = 3, f = 15, g = 3, h = 21, i = 9). (E) Dimensionality reduction and cluster validation. Left: t-SNE embedding (perplexity = 30) of genera colored by cluster; gray ellipses denote 95% confidence regions; dashed arrow indicates pseudo-temporal gradient. Right: distance-to-centroid distributions (violins) demonstrating cluster compactness (PERMANOVA p < 0.01). (F) Hierarchical heatmap of 77 significant genera (Kruskal–Wallis, FDR < 0.05) with Ward’s linkage dendrograms. Rows annotated by phylum and cluster; columns ordered by age group; color scale shows standardized abundance. (G) Distributions of Shannon (Top) and Bray–Curtis (Bottom) diversity metrics per age-group. Red line indicates mean values, Kruskal–Wallis FDR P-value, R-squared and sample size shown.

**Table 1:** Summary of microbiome wide association model specifications in this study.

| Analysis Type | Dependent Variables (M) | Predictors per Dep. Variable (N) | Covariates | Total Tests | p<0.05 | p<0.01 | FDR p<0.05 | FDR p<0.01 | Q-value< 0.05 | Q-value< 0.01 |
| --- | --- | --- | --- | --- | --- | --- | --- | --- | --- | --- |
| Demographics → Microbiome (Linear) | 1,349 OTUs | 13 demographics | None | 17,537 | 11,493 (65.5%) | 9,775 (55.7%) | 11,055 (63.0%) | 8,443 (48.1%) | 13,742 (78.4%) | 11,425 (65.2%) |
| Microbiome → Oral Conditions (Logistic) | 4 traits | 1,349 OTUs | 13 demographics | 5,396 | 3,893 (72.2%) | 2,910 (53.9%) | 3,388 (62.8%) | 2,672 (49.5%) | 5,288 (98.0%) | 4,326 (80.2%) |
| Blood/Urine Markers → Microbiome (Linear) | 1,349 OTUs | 473 exposures | 13 demographics | 638,077 | 95,122 (14.9%) | 42,970 (6.7%) | 22,108 (3.5%) | 12,425 (1.9%) | 25,429 (4.0%) | 12,788 (2.0%) |
| Microbiome → Measured Phenotypes (Linear) | 133 phenotypes | 1,349 OTUs | 13 demographics | 179,417 | 15,935 (8.9%) | 4,985 (2.8%) | 172 (0.1%) | 12 (0.0%) | 327 (0.2%) | 33 (0.0%) |
| Microbiome → Disease Incidents (Logistic) | 16 outcomes | 1,349 OTUs | 13 demographics | 21,584 | 5,394 (25.0%) | 1,876 (8.7%) | 845 (3.9%) | 285 (1.3%) | 971 (4.5%) | 325 (1.5%) |

Across demographic variables, many oral genera showed significant associations in abundance (FDR < 0.05) (Figure 1B). *Filifactor, Aggregatibacter, Peptococcus*, and *Peptostreptococcus* abundance was shown to be positively correlated with Mexican and other Hispanic ethnicities compared with White individuals, while Black participants were associated with higher levels of *Anaeroglobus*, *Filifactor*, and *Lactobacillus*. *Proteobacteria* phyla such as *Actinobacillus*, *Neisseria*, *Mannheimia*, and *Lautropia* were significantly differentially abundant in Mexican and other Hispanic participants, whereas several *Firmicutes* and *Actinobacteria*—including *Bifidobacterium*, *Alloscardovia*, *Atopobium, Megasphaera*, and *Scardovia*—were more abundant in White participants (Fig 1C). Education-related associations were modest, with *Cardiobacterium* abundant among individuals with higher educational attainment. *Peptoclostridium* and other anaerobic taxa were associated with lower education levels (Figure 1B and 1C). We also note that unlike the previous study findings in oral microbiome^24^, Shannon alpha diversity showed highest in Black and lowest in White NHANES study participants (see Supplementary Tables S5 and S7). All other groups also showed higher Shannon diversity than White: Mexican American, Other, and Other Hispanic. Bray–Curtis beta diversity differed significantly from White for Mexican American, Other, and Other Hispanic, however Black population showed no significant separation from White population (see Supplementary Tables S5 and S6).

We call the set of regression estimates between a host variable and all tested bacterial taxa, summarized by its taxon-level regression estimate a “signature”. To capture the full association between total microbial community composition and host phenotype, we calculated inverse-variance weighted correlations between the computed microbial signatures and hierarchically clustered these relationships (Fig 1D; see *Methods*). We observed expected correlations structures, or signatures; for example, “Mexican American” and “Other Hispanic” identities shared a closely aligned microbiome signature, whereas gender and age related variables formed distinct clusters, indicating a characteristic signature largely independent of other demographic traits.

We next aimed to identify network structures of the demographically associated taxa (as observed in Fig 1D). We constructed association networks that capture the direction and magnitude of each taxon’s effect and visualize how these effects co-vary across the community.^25^ In the gender-associated network (Figure 1E, left), female-enriched taxa were dominated by *Proteobacteria* (*Pseudomonadota*)—including *Pseudomonas*, *Azomonas*, *Paracoccus*, and *Burkholderia*—which are non-spore-forming, motile, rod-shaped, and largely aerobic. Male-associated taxa instead harbored a wider variety, mainly *Firmicutes* and *Actinobacteria*, such as *Filifactor*, *Peptococcus*, *Peptoclostridium*, *Megasphaera*, and *Bifidobacterium*.

Beyond testing genus-level associations with host features, we integrated microbial phenotype annotations from the JGI-GOLD database to compute phenotype indices (gram stain, motility, oxygen requirement, and spore-formation capacity; see Methods). Notably, this approach enables quantification of validated functional traits for each identified genus and systematic aggregation of these traits among microbes significantly associated with variables of interest. For each host variable, we present the distribution and mean of microbial phenotype scores for taxa with positive versus negative associations, allowing a community-level ecological interpretation rather than focusing on individual taxa.

### Heterogeneous, Non-Linear Age Trajectories Across Oral Genera

The oral microbiome was highly indicative of host age. 118 unique genera (out of 1,349 tested) exhibited significant linear or quadratic associations with age (FDR < 0.05, prevalence > 10%), including 108 with linear age associations (63 with complete annotations) and 111 OTUs with quadratic age associations (64 with complete annotations) (Figure 2A; Supplementary Table S2 for full result). These included *Alloscardovia*, *Bifidobacterium*, *Rothia*, and *Prevotella* in older adults and *Streptobacillus*, *Actinobacillus*, *Aggregatibacter*, and *Curvibacter* in younger individuals (Fig. 2B). In general, we found that inter-individual β-diversity (e.g., Bray–Curtis dissimilarity) rose steadily across the lifespan, whereas α-diversity peaked in young adulthood (20–29 years) before declining (Figure 2G).

Next, we hypothesized that additional latent age trajectories (e.g., segments of the population that exhibit a specific “type” of correlation with age, or “age-o-type”^26^) might exist beyond the population-based trends identified in the linear regression. Among prevent (>10% prevalence) genera (N = 93), we observed a wide range of age-related abundance patterns, from approximately linear shifts to complex non-linear trajectories (Figure 2D). To characterize the underlying patterns, we analyzed standardized (z-score) mean abundances of these 93 genera across eleven age groups spanning 14–19 to 65–69 years. We identified nine optimal clusters (k = 9; silhouette = 2.851) of patterns using “Silhouette maximization”. Each latent cluster represented an age-related trajectory (Figure 2C and 2D). Cluster sizes ranged from 3 to 21 genera, indicating diversity in age-dependent dynamics among common members of the oral microbiome.

We used dimensionality reduction to evaluate separation among latent trajectories. The resulting clustered embedding (Fig. 2E) showed compact, non-overlapping cluster neighborhoods arranged along an age gradient from adolescence to late adulthood (dashed arrow). Distance-to-centroid distributions indicated significant separation between clusters (PERMANOVA p < 0.01), and cluster identities remained stable across both PCA and t-SNE embeddings, demonstrating internal consistency of such pattern (Supplementary Figure S3). We interpret this ordering as reflecting cluster-specific ages at which genera reach peak abundance.

We highlight the diagonal gradients that capture shifts in relative abundance (77 genera), delineating potential genus-specific accumulation modules (Fig. 2D and F). Early-peaking clusters (e.g., a–c) were enriched for genera such as *Curvibacter*, *Haemophilus*, *Corynebacterium*, *Alloprevotella*, and *Streptobacillus*, whereas mid-life–peaking clusters (e.g., d–f) were characterized by *Dialister*, *Campylobacter*, *Prevotella*, *Treponema*, and *Fusobacterium*. In contrast, late-increasing clusters (e.g., h, i) included genera such as *Rothia*, *Leptotrichia*, *Veillonella*, *Alloscardovia*, and *Bifidobacterium*, indicating coordinated, phylum-level transitions from early *Firmicutes*/*Bacteroidetes* dominance toward *Actinobacteria*-rich communities in later adulthood.

### Periodontal Pathobionts are associated with Gum Disease, Poor Oral Hygiene, and Dental Caries

We examined associations between microbial composition and four local binary oral conditions, including gum disease, oral hygiene, tooth decay, and denture use. These diagnoses arose from questionnaires, for example, participants were asked, “has a dentist ever diagnosed you with gum disease?”. Gum disease, poor oral hygiene, and tooth decay were each associated with gram-positive genera and were predominantly anaerobic species (Figure 3A). Across the four conditions, poor oral hygiene displayed a microbial signature (Figure 3B) most similar to that of gum disease, with substantial overlap with tooth decay, whereas denture use exhibited a distinct correlation structure as depicted by the network diagrams. When restricting the analysis to the most prevalent taxa and their associations with oral conditions (Figure 3C), we found that the denture environment represents a distinct niche relative to the other conditions.

**Figure 3.**
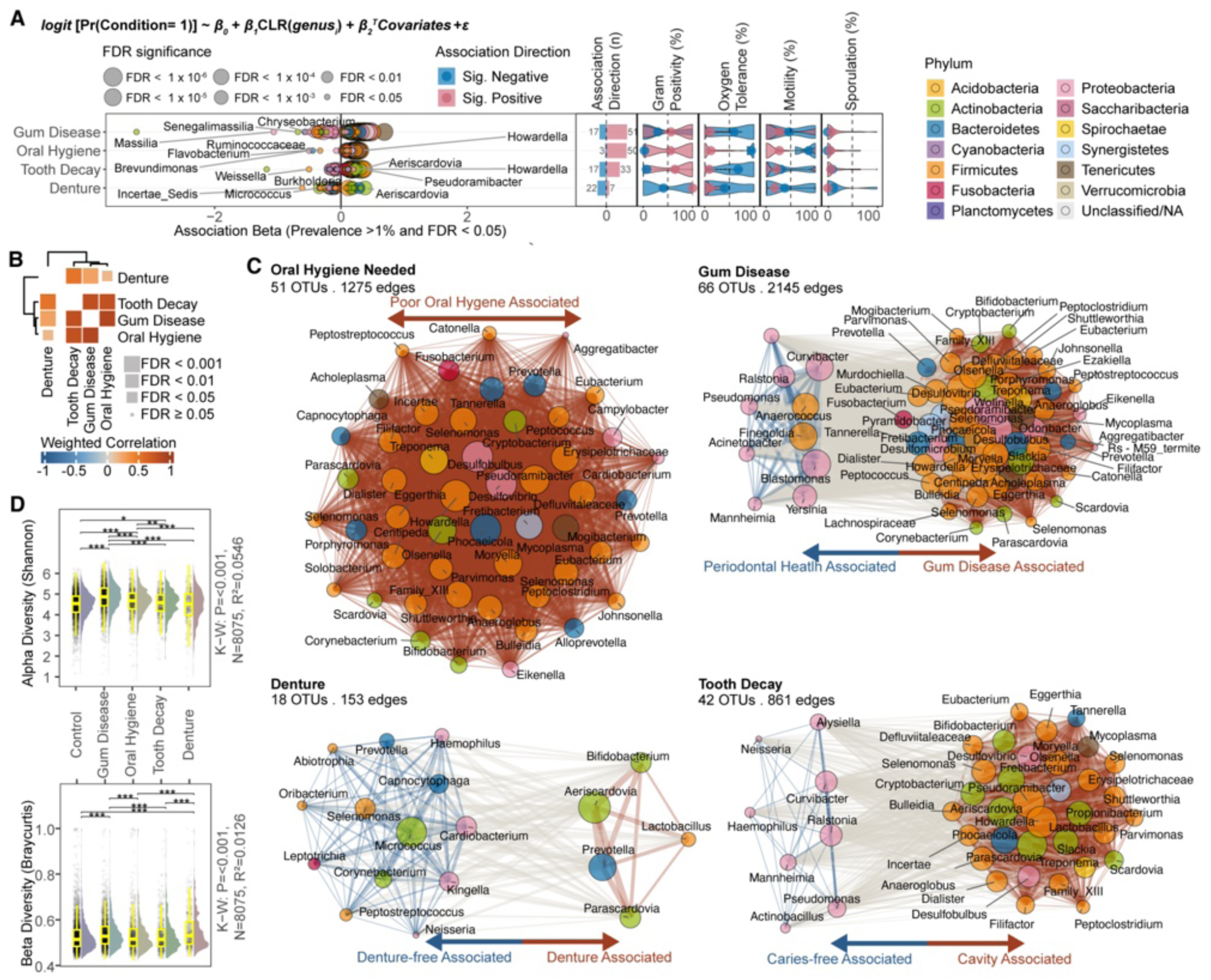
Oral microbiome association with local conditions in the oral cavity. Denture exhibits a distinct relationship architecture. (A) Survey-weighted logistic regression between 1,349 OTUs and four local oral conditions across NHANES 2009–2010 and 2011–2012 (FDR ≤ 0.05, q ≤ 0.05; prevalence ≥ 1%) adjusted for host demographics. Each point represents an effect size (x-axis), with dot size denoting FDR category and color indicating phylum. Right panels summarize significant associations by direction (positive/negative) and predefined microbial indices. (B) Weighted correlation matrix of microbial association signatures across the four oral conditions, showing FDR threshold (square size) and correlation strength (color). (C) Small effect network of oral-condition versus control group oral genera associations (FDR ≤ 0.05, q ≤ 0.05; prevalence ≥ 1%). Nodes represent genera, with size proportional to cumulative effect magnitude and color indicating phylum. Edge width reflects geometric mean of effect sizes, and color encodes directional concordance (orange = positive; blue = negative). (D) Alpha (Shannon; top) and beta (Bray–Curtis; bottom) diversity distributions with pairwise Wilcoxon comparisons; brackets indicate significance (*** FDR < 0.001, ** FDR < 0.01, * FDR < 0.05).

Numerous genera associated with gum disease and poor oral hygiene were aligned with well-established periodontal pathogens and dysbiotic community members. In particular, our results (Figures 3B and 3C) shows that *Campylobacter* (in poor oral hygiene), *Fusobacterium*, *Porphyromonas*, *Prevotella*, *Peptostreptococcus*, *Tannerella*, and *Treponema* were all significantly positively associated to these conditions, consistent with decades of subgingival plaque and periodontitis research.^27–29^ In addition, our findings (Figure 2C) also supporting significant associations of *Aggregatibacter*, *Eubacterium*, *Parvimonas*, and *Prevotella* tightly track the broader periodontal health field’s knowledge^30,31^.

We found that caries-free and periodontally healthy subjects exhibited significantly higher abundances of numerous *Proteobacteria*, including *Curvibacter*, *Ralstonia*, *Pseudomonas*, *Actinobacillus*, and *Mannheimia*—a pattern consistent with prior reports identifying *Proteobacteria* as characteristic of “healthy” oral microbiota^29^. We note that among *Proteobacteria* phyla found to be weakly negatively associated with dental caries, *Actinobacillus* (*Aggregatibacter*), a well-characterized cavity- and periodontitis-associated oral pathogen^32,33^, also showed a negative association. Healthy oral conditions (No Cavities, Healthy Gums) were generally dominated by aerobic and facultative taxa, reflecting a well-oxygenated and disrupted plaque environment. Among these, *Neisseria* and *Haemophilus*—canonical early colonizers enriched in periodontal health^34,35^ were among the strongest markers of the healthy group, consistent with previous observations^28^. Additional oxygen-tolerant genera such as *Alysiella*, *Aggregatibacter*, *Ralstonia*, *Pseudomonas*, and *Blastomonas* were likewise enriched, supporting a community structured around oxygen-utilizing, non-proteolytic microbes.

Commensal gram-positive anaerobes including *Anaerococcus* and *Finegoldia*, hypothesized to be mucosal residents^36^ were also significantly associated with healthy gums, likely reflecting the reduced presence of disease-associated anaerobes rather than a direct protective function. In contrast, classical early colonizers such as *Streptococcus* and *Actinomyces* did not distinguish health status, consistent with their broad distribution across oral conditions^29,37^. Collectively, these patterns delineate a *Proteobacteria*-rich, aerobes-favored community characteristic of low-inflammation oral health.

### Microbial Signatures of Local Oral Conditions Reveal a Distinct Denture-Associated Niche

Our analysis shows that denture wearers consistently exhibit an aciduric, cariogenic microbial profile—associations dominated by *Lactobacillus*, *Bifidobacterium*, *Scardovia*/*Parascardovia* and *Aeriscardovia* (Figure 3A and 3C)^38^. Denture surfaces create a low-pH, plaque-like niche favoring these taxa along with mutans *streptococci* and yeasts^39^, compared to healthy dentate individuals. B*ifidobacteria* can persist in denture environments^39^, consistent with our findings. Our observation of *Prevotella* associations appears to be context-dependent: while *Prevotella* is reported to be reduced in edentulous conditions^40^, it has been shown to persist in partial denture wearers^38^ and to increase under inflammatory or recently re-colonized denture conditions^41,42^. We report that genera negatively associated with denture wearers were *Haemophilus*, *Neisseria*, *Corynebacterium*, *Selenomonas*, *Leptotrichia*, *Capnocytophaga*, and other tooth-associated commensals, that supports our understanding that edentulism eliminates key ecological niches required for their persistence and introduces cleaning regimens specific to dentures^38,43^. We also identified several less-reported genera that are negatively associated with denture wear, including *Peptostreptococcus*, *Abiotrophia*, *Oribacterium*, *Kingella*, and *Cardiobacterium*, which likely reflect their adaptation to natural tooth architecture.

### Oral microbiome composition is correlated to smoking and its associated biomarkers

Across the 473 serum, blood, and questionnaire exposure markers, smoking- and combustion-related biomarkers stood out immediately in association with the oral microbe genera. Each of these markers were significantly associated with more than 50 oral genera (Figure 5A). Prior studies of the American oral microbiome^44^ report lower abundance of *Capnocytophaga*, *Peptostreptococcus*, *Leptotrichia*, and multiple *Proteobacteria* species in smokers, alongside higher levels of *Atopobium* and *Streptococcus* genera.

Our result shows that many exposure variables likely linked to smoking exhibited highly similar microbiome signatures (Supplementary Figure S8). Biomarkers such as blood 2/3-fluorene, blood 2,5-dimethylfuran, blood furan, and blood benzene clustered closely with one another and showed association patterns that closely mirrored those patterns associated with participant smoking status (Supplementary Figure S8).

Circulating cotinine (a metabolite of nicotine) levels were negatively associated with several aerobic and aerotolerant genera, including *Neisseria*, *Haemophilus*, *Gemella*, *Lautropia*, *Actinobacillus* (*Aggregatibacter*), and *Cardiobacterium* within the *Proteobacteria* phylum, as well as *Capnocytophaga*. In contrast, taxa showing positive associations with cotinine were dominated by obligate anaerobic and acid-tolerant genera. These included *Lactobacillus* and *Bifidobacterium*, as well as *Scardovia*-related genera such as *Alloscardovia*, *Parascardovia* and *Metascardovia* all of which we also reported to be associated with tooth decay and gum disease (Figure 3C). Cotinine was also positively and significantly associated with several anaerobic genera characteristic of low-oxygen or inflammatory oral niches, including *Treponema*, *Pseudoramibacter*, *Fretibacterium*, *Filifactor*, *Bulleidia, Cryptobacterium*, *Mycoplasma*, *Anaeroglobus*, *Howardella*, *Olsenella*, and *Phocaeicola*. Collectively, these cotinine-associated taxa reflect a shift toward aciduric and anaerobic metabolism at the expense of normal oxygen-tolerant commensals.

However, amid these shared signatures across biomarkers of smoking exposure biomarkers (e.g., cotinine)(Figure 4A), we highlight clear points of divergence, reflecting their distinct exposure profiles (Figure 4C). Cotinine is a metabolite of nicotine and will reflect recent nicotine intake or second hand exposure, and was associated with a broader depletion of oxygen-tolerant commensals. In contrast, cadmium, which captures long-term accumulation of tobacco-derived heavy metals, showed a narrower but distinct set of FDR-significant negative associations, uniquely with *Eikenella*, *Moraxella*, *Alysiella*, and *Johnsonella*. Cadmium’s positive associations closely mirrored those of cotinine, with no additional cadmium-specific enriched taxa beyond this shared anaerobic group. Cadmium is found most prominently in cigarette smoke, although it is also found in food^45^.

**Figure 4.**
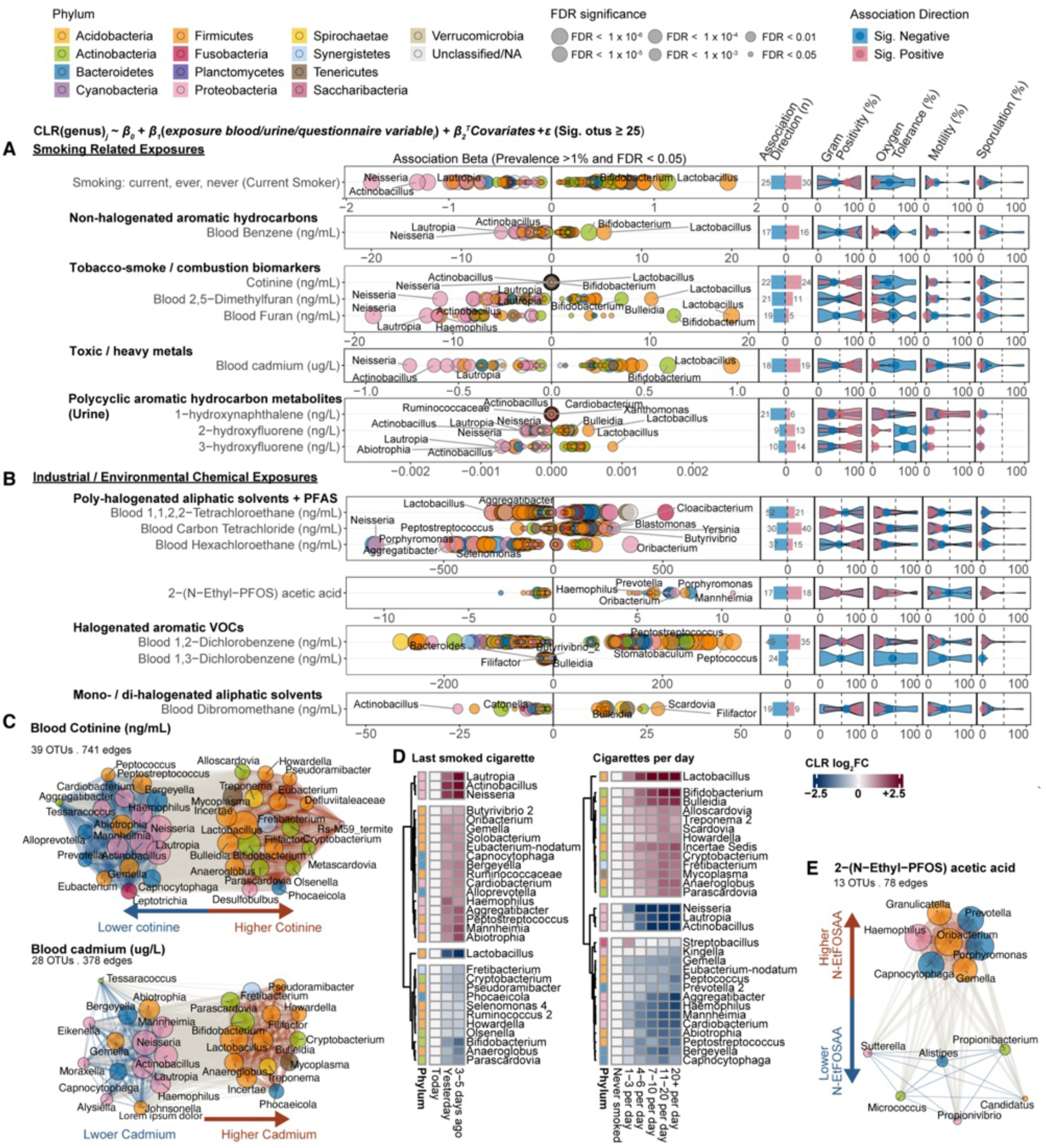
Smoking-related exposure signatures in the oral microbiome across biomarker, questionnaire, and ecological context. (A) Survey-weighted regressions (NHANES 2009–2010, 2011–2012) of smoking-related variables with oral genera, restricted to exposures with ≥25 significant associations (β₁; prevalence ≥1%, FDR ≤0.05). Left: regression coefficients; dot size indicates FDR significance. Right: GOLD-DB microbial phenotype indices (Gram positivity, oxygen tolerance, motility, sporulation) summarized for annotated genera by association direction. (B) Same regression layout as in (A), applied to additional chemical exposure variables. (C) Small-effect network of significant and common genus–exposure associations for representative smoking biomarkers (prevalence ≥10%, FDR ≤0.05). Node size reflects cumulative effect magnitude; node color denotes phylum; edge color indicates association direction (blue, negative; orange, positive). (D) Heatmaps of mean CLR abundances across smoking-intensity and smoking-recency categories. Cell colors represent CLR log₂ fold-change (capped at ±2.5). Rows show the 30 most variable genera (prevalence ≥1%, non-zero CLR variance, FDR ≤0.05), identified using Kruskal–Wallis tests and clustered by Euclidean distance. (E) Small-effect network (as in C) for 2-(N-ethyl-PFOS) acetic acid.

**Figure 5.**
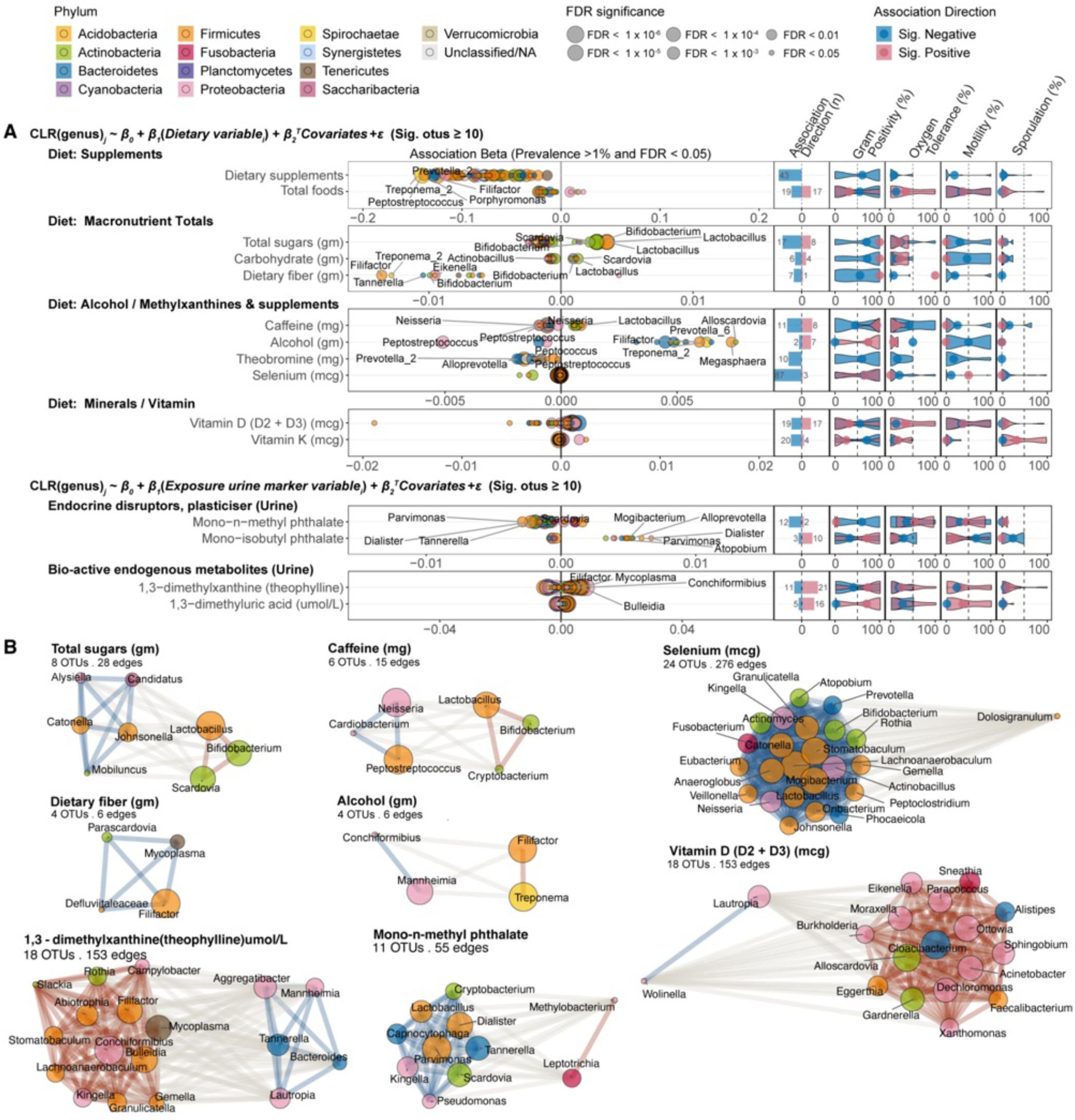
Dietary and other exposures correlates with the residents of the oral microbiome. (A) Survey-weighted association results across two NHANES oral microbiome cycles (2009–2010, 2011–2012) with reported dietary and other urine marker variables; model specification as indicated and rows are restricted to variables with at least 10 significant (β1; prevalence ≥ 1%, FDR ≤ 0.05, q ≤ 0.05) oral microbes. Left: regression coefficients (β1) with dot size indicating FDR significance. Right: GOLD-DB–derived microbial trait indices (Gram positivity, oxygen tolerance, motility, sporulation) summarized by association direction (positive/negative). Only variables with non-NA GOLD-DB–annotated traits are displayed. (B) Selected networks of small effect ecology (FDR ≤ 0.05; prevalence ≥ 10%) with dietary variables. Nodes represent genera, size showing cumulative effect magnitude and color indicating phylum, edge color encodes direction (blue = negative, orange = positive). (C) Association results for urine marker variables; specifications as indicated. (D) Small-effect networks for the selected variables related to urine exposure marker variable (see Supplementary Figures S9 and S10 for full list).

We reasoned that these broadly responsive genera might also reflect dose-dependent and recency-dependent patterns of tobacco exposure (Figure 4D). Across both dimensions of exposure, we observed a highly coherent pattern: anaerobic and acidogenic genera—such as *Lactobacillus*, *Bifidobacterium*, *Scardovia*/*Alloscardovia*, *Parascardovia*, *Treponema*, *Fretibacterium*, *Cryptobacterium*, *Howardella*, and *Anaeroglobus*—rose steadily with increasing cigarettes smoked per day and showed the opposite trend with greater time since last cigarette, indicating rapid suppression during abstinence. In contrast, oxygen-tolerant commensals including *Neisseria*, *Lautropia*, *Actinobacillus*, *Aggregatibacter*, *Haemophilus*, *Gemella*, *Capnocytophaga*, *Streptobacillus*, *Kingella*, *Cardiobacterium*, *Bergeyella*, *Abiotrophia*, *Peptostreptococcus*, *Mannheimia*, and *Eubacterium* nodatum displayed reciprocal patterns, declining with heavier smoking and rebounding within days of cessation. Superimposed on this shared axis were a small number of exposure-specific taxa: *Bulleidia*, *Peptococcus*, and certain *Prevotella* showed clear dose-dependent responses without short-term recency effects, whereas *Butyrivibrio*, *Oribacterium*, *Solobacterium*, *Ruminococcaceae*, *Alloprevotella*, *Pseudoramibacter*, *Phocaeicola*, *Olsenella*, *Selenomonas*, and related genera were uniquely responsive to short abstinence intervals.

### Chemical and environmental toxicants exhibit associations with the oral microbiome

A similar oral-microbiome association structure emerged when we turned from behavioral exposures to chemical toxicants. Among a distinct set of exposures, circulating chlorinated solvents—including 1,1,2,2-tetrachloroethane, carbon tetrachloride, and hexachloroethane—showed broad yet taxonomically heterogeneous associations with the oral microbiome (Figure 4B). We additionally identified 2-(N-ethyl-perfluorooctane-sulfonamido) acetic acid (a downstream product of the perfluorinated surfactant PFOS) as exhibiting a highly skewed profile, associating positively with *Prevotella*, *Porphyromonas*, *Capnocytophaga*, and other anaerobes. We likewise observed that several halogenated aromatic VOCs (blood 1,2-dichlorobenzene and 1,3-dichlorobenzene) and mono-/di-halogenated aliphatic solvents (e.g., blood dibromomethane) were associated with numerous oral taxa in both positive and negative directions. However, given the broad chemical reactivity, mixed exposure sources, and relatively low biomarker concentrations for these compounds, these patterns warrant cautious interpretation (Supplementary Figures S6 and S7).

### Dietary Intake Influences Oral Community Structure

We further examined associations between reported dietary intake variables and oral microbial abundances (Figure 5). We show that the total number of dietary supplements reported within the 24-hour recall was broadly negatively associated with many predominantly anaerobic oral genera, substantially overlapping with taxa negatively associated with selenium intake (Figure 5A and 5B; Supplementary Figure S9 and S10). We also note that the total number of foods reported was negatively associated with a wide range of phyla and genera, including *Johnsonella*, *Phocaeicola*, *Fretibacterium*, and *Filifactor*, and positively associated with oxygen-tolerant taxa such as the *Proteobacteria* genus *Simonsiella*, though it only captures a limited portion of the overall relationship between diet and oral environment.

Building on our observation that acidogenic, cariogenic taxa—including *Lactobacillus*, *Bifidobacterium*, and *Scardovia* (Figure 3)—were positively associated with disease, we found that these contributors to dental caries^46,47^ were also the genera most positively correlated with reported sugar, carbohydrate, and caffeine intake (Figure 5A and 5B; Supplementary Figure S10).

We note that alcohol intake shifted the oral microbiome toward a periodontal-disease–like state, marked by increased *Filifactor* and *Treponema*^48^. Also, anaerobic commensals such as *Alysiella*, *Catonella*, *Johnsonella*, and *Mobiluncus* were negatively associated with sugar intake, well-supported by the ecological plaque hypothesis of suppressed acid-intolerant taxa^49^. Although the caffeine metabolite theophylline showed negative associations with genera such as *Aggregatibacter*, *Mannheimia*, *Tannerella*, *Bacteroides*, and *Lautropia*, and positive associations with a broader and more heterogeneous set including *Rothia*, *Gemella*, *Granulicatella*, *Slackia*, *Bulleidia*, *Filifactor*, and *Mycoplasma*, these patterns diverged from those observed for caffeine intake itself (Figure 5A and 5B). Rather than reflecting a direct metabolic effect, this signature likely warrants more cautious interpretation and may be better explained by smoking-related microbial shifts, as explored in Figure 4. We also report that higher dietary fiber intake showed broadly negative associations with genera linked to oral disease—including *Parascardovia*, *Filifactor*, *Mycoplasma*, and a *Defluviitaleaceae* genus—none of which have previously been reported to decrease in dietary intervention studies involving higher fiber intake^50,51^. Together, these suggest that excess sugar and carbohydrate promotes cariogenic and aciduric taxa, whereas fiber-rich diets suppress pathogenic oral taxa without selectively enriching specific beneficial genera.

We found that selenium showed consistently negative correlations with many oral bacteria, particularly *Firmicutes* and *Proteobacteria*. This is notable because selenium exposure has been shown to alter gut microbiome composition in mouse models by modulating antioxidant and inflammatory pathways^52^, and dietary selenium is suggested to interact bidirectionally with the gut microbiota—both affecting selenium bioavailability and influencing microbial communities through host selenoprotein regulation^53,54^. Given longstanding evidence dating back to the 1960s that selenium can influence enamel surface properties and has been linked to negative effects on dental caries and oral health^55–57^.

Vitamin D intake showed mixed association directions, with several mostly environmental and anaerobic genera (including *Paracoccus*, *Ottowia*, *Cloacibacterium*, *Acinetobacter*, *Dechloromonas*, *Xanthomonas*, and *Burkholderia*) increasing, while taxa such as *Wolinella* decreased. Vitamin K intake was associated with decreases in multiple anaerobic, periodontal-associated genera (e.g., *Porphyromonas*, *Peptostreptococcus*, *Solobacterium*; see Supplementary Figures S9, S10 and Table S2 for full results). We also note that urinary Mono-nmethyl Phthalate (MMP), a phthalate metabolite, was associated with broad decreases in several oral genera (including *Cryptobacterium*, *Lactobacillus*, *Dialister*, *Capnocytophaga*, *Tannerella*, *Parvimonas*, *Kingella*, *Scardovia*, and *Pseudomonas*), alongside increases in *Methylobacterium* and *Leptotrichia*. Although prior work^58^ suggested that multiple phthalate metabolites (e.g., MECPP, MnBP, MEHHP, MEOHP, MBzP) are linked to higher periodontitis risk, this pattern remains difficult to interpret.

### Oral microbial signatures extend to systemic health and broader physiomic states

To extend our findings that oral microbial shifts are linked to local oral conditions and to multiple environmental and chemical exposures, we next asked whether these microbial variations also relate to broader systemic health. Because oral communities interface continuously with host immunity, circulation, and inflammation, as a natural progression of our investigation, we evaluated whether individual oral genera were associated with (i) clinically diagnosed systemic conditions, (ii) quantitative host phenotypes, and (iii) immune marker profiles (Figure 6). Using fully adjusted logistic and linear models and focusing on outcomes with at least five FDR-significant genera, we identified specific diseases, metabolic traits, and immune indicators that show robust oral-microbiome associations (Figure 6A). This framework allows us to move from exposure-linked and locally confined oral effects toward a broader characterization of how oral ecology affects host physiome.

**Figure 6.**
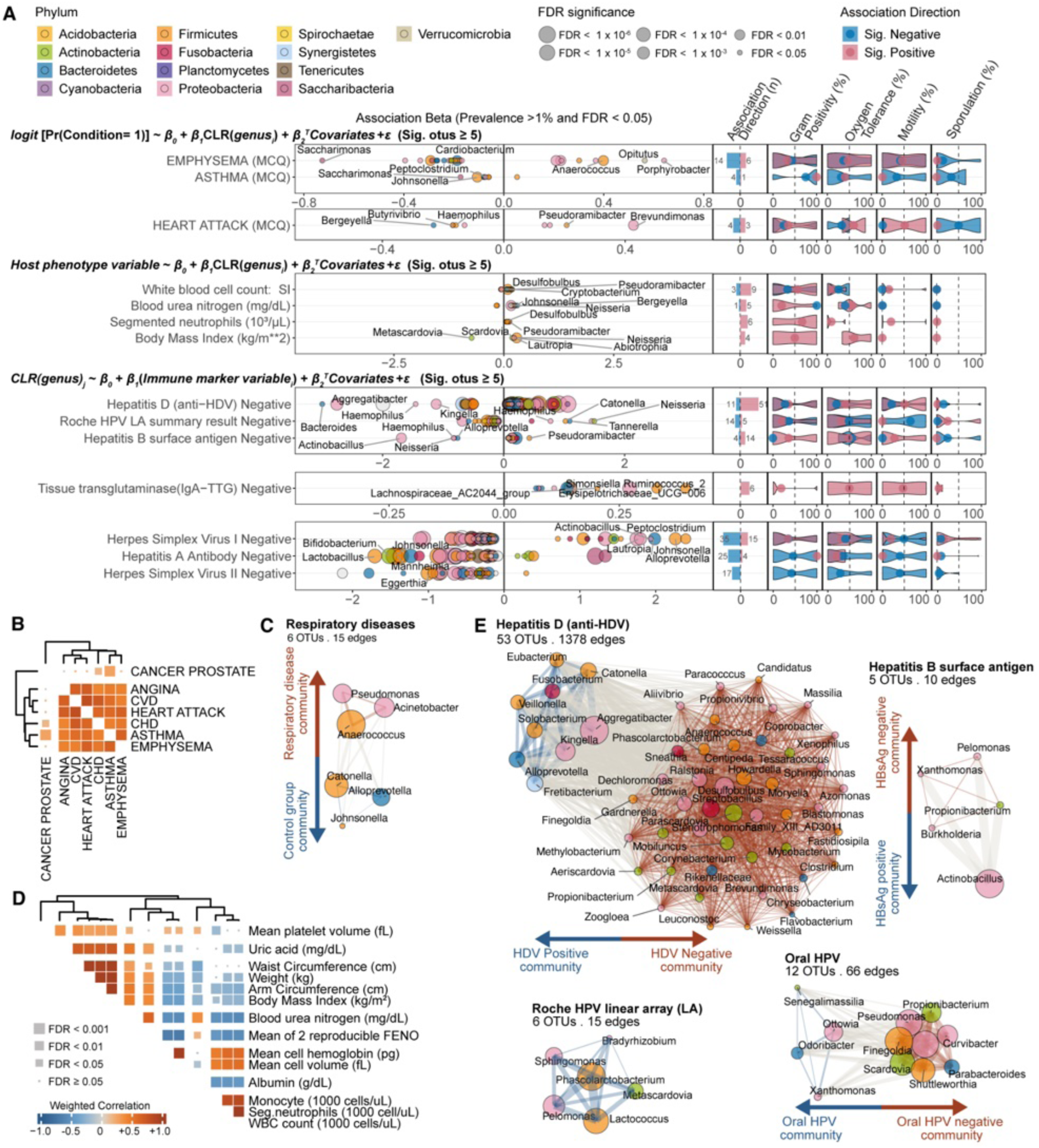
Oral microbial genera is more associated with immunological markers than measurable host phenotypes and outcome measures. Survey-weighted regressions across NHANES oral microbiome cycles (2009–2010, 2011–2012). (A) Associations with reported non-oral disease incidents, measurable phenotypes and blood immune marker variables are shown with the model specification as indicated; restricted to variables with at least 5 significant (β1; prevalence ≥ 1%, FDR ≤ 0.05, q ≤ 0.05) oral microbes. Left: regression coefficients (β1) with dot size indicating FDR significance. Right: GOLD-DB–derived microbial phenotype indices (Gram positivity, oxygen tolerance, motility, sporulation) summarized by association direction (positive/negative). Only variables containing at least one significant and GOLD-DB–annotated genus are displayed. (B) Microbial association signatures across reported disease incidents. Restricted to variables with at least two significantly associated (FDR < 0.05) common microbes (prevalence > 10%). Each cell represents the inverse-variance–weighted Pearson correlation between pairs of (β) estimates derived from CLR-transformed models and cell size indicates FDR significance. (C) Grouped small effects network of all reported respiratory disease incidents (see Methods). (D) Microbial association signatures across all measurable phenotype variables. (E) Four selected small effects networks of oral ecology (FDR ≤ 0.05, q ≤ 0.05; prevalence ≥ 1%). Nodes represent genera, with size proportional to cumulative effect magnitude and color indicating phylum.

### Respiratory and Cardiovascular Links to Oral Taxa Reflect Core Respiratory-Pathogen Biology

Across the 16 participants-reported diseases, emphysema, asthma, and heart attack demonstrated the strongest oral-microbiome associations, each involving at least five genera that reached FDR significance. Disease-level microbial signature analyses further revealed that angina, cardiovascular disease (CVD), and heart attack shared highly similar microbial profiles, whereas asthma and emphysema, a subtype of chronic obstructive pulmonary disease (COPD) formed a distinct cluster characterized by shared association patterns across all 1,349 genera tested (Fig. 6B). Given that the oral–lung axis constitutes a shared ecological environment in which oral and lung microbial communities are tightly interconnected and jointly influence lung and airway disease etiology, this clustering pattern is biologically plausible rather than trivial. We find that disease groups in our dataset, including emphysema and asthma showed statistically significant α- and β-diversity differences relative to controls (Supplementary Tables S5, S6 and S7). Consistent with our diversity findings, earlier NHANES-based COPD work^59^, reported reduced α-diversity and corresponding β-diversity differences in COPD. However, these differences were uniformly small in magnitude, suggesting limited biological interpretability. Accordingly, we cautiously reason that the microbiome signatures observed here are driven primarily by taxon-specific shifts rather than broad changes in overall diversity–disease relationships.

Heart attack was associated with lower abundance of *Haemophilus*, *Bergeyella*, and *Butyrivibrio*, alongside higher abundance of *Pseudoramibacter* and *Brevundimonas* (Fig. 6A). Emphysema showed broad inverse associations with multiple anaerobic genera (*Alloprevotella*, *Saccharimonas*, *Cardiobacterium*, *Catonella*, *Johnsonella*), with positive associations involving *Anaerococcus*, *Opitutus*, and several *Proteobacteria*, including *Porphyrobacter*, *Pseudomonas*, and *Acinetobacter*. Asthma demonstrated a similar but less extensive pattern, marked by reduced abundance of *Saccharimonas*,

*Peptoclostridium*, and *Johnsonella*. To identify microbial signals that are extensively shared across respiratory conditions, we aggregated associations for asthma, emphysema, and bronchitis. The resulting grouped respiratory-disease association network (Fig. 6C) highlights taxa meeting the ≥10% prevalence threshold and demonstrating at least two directionally concordant associations. This compact set of genera reveals that respiratory disease is most consistently and positively associated with *Pseudomonas*, *Acinetobacter*, and *Anaerococcus*, while negatively associated with *Catonella*, *Alloprevotella*, and *Johnsonella*.

This pattern aligns with recent evidence showing that *Acinetobacter*—a taxon, when residing in respiratory tract, implicated in smoking-related reductions in lung function and enriched metabolic, resistance, and virulence-factor pathways^60^ and *Pseudomonas*, widely recognized for potent virulence factors and its central role in chronic respiratory infections^61^ are both strongly linked to adverse respiratory outcomes. The well-established respiratory pathogenicity of *Pseudomonas* is further supported by studies showing pyocyanin-mediated epithelial damage(<u>Rada & Leto, 2013</u>), highly adaptable aerobic and anaerobic respiratory metabolism enabling airway persistence^62^, chronic pulmonary infection in the classic rat agar-bead model^63^, and adhesion to respiratory mucins via defined adhesin–receptor interactions^64^. Although the positive association between respiratory disease and the *Anaerococcus* genus remains understudied, our results support a recent case report of a rare *Anaerococcus prevotii*–driven pulmonary infection^65^, which describes a severe pleural empyema with extensive suppuration caused by *A*. *prevotii* together with *Fusobacterium nucleatum*.

### Inflammatory and Metabolic Traits Associate with Distinct Oral Microbial Signatures

Across 133 phenotype measures, four traits, white blood cell (WBC) count, segmented neutrophils count, blood urea nitrogen (BUN), and BMI, each associated with multiple oral genera (Figure 6A, middle). Elevated BUN correlated with increased abundances of *Johnsonella*, *Bergeyella*, and *Neisseria*, taxa not previously linked to BUN. Higher BMI was associated with *Lautropia*, *Neisseria*, and *Abiotrophia*, and inversely associated with *Metascardovia*. These patterns suggest possible oral–renal and oral–metabolic connections, though further investigation is necessary. Inflammatory cell counts (WBC, segmented neutrophils, and monocytes) shared strongly overlapping microbial correlates (Figure 6D), indicating a common inflammation-linked oral signature. In contrast, adiposity-related traits (BMI, waist circumference) displayed microbiome patterns more similar to those of uric acid and mean platelet volume than to leukocyte measures, suggesting a distinct metabolic–rather than inflammatory–microbial profile.

WBC and segmented neutrophils were positively associated with higher CR abundance of *Desulfobulbus*— sulfate-reducing periodontal pathobiont genera such as *D. orale*^66^ isolated from deep oral sites and repeatedly linked to periodontitis^66–68^ and hyperglycemia-associated^69^ dysbiosis—as well as *Pseudoramibacter*, a dominant anaerobe in infected root canals^70,71^, and *Cryptobacterium*, a Gram-positive anaerobe found in periodontal pockets^72^ and other endodontic infections. Collectively, these taxa are consistently enriched in periodontal or endodontic disease across studies, suggesting that the elevated leukocyte-linked microbial signatures in our cohort might be reflective of increased oral anaerobic burden and subclinical oral inflammation rather than nonspecific systemic inflammation.

Serologic markers of viral infection, including hepatitis D virus (HDV), human papillomavirus (HPV), hepatitis B surface antigen (HBsAg; HBV), herpes simplex viruses (HSV-1/2), and hepatitis A, showed broader associations with oral taxa than host phenotypes or other disease-related variables (Fig. 6A). We show that participants positive for general HPV had enrichment of uncommon genera such as *Lactococcus*, *Pelomonas*, *Phascolarctobacterium*, *Sphingomonas*, and *Metascardovia* (Fig. 6A, 6E), whereas oral HPV positivity was instead associated with increased *Ottowia*, *Odoribacter*, and *Xanthomonas*, indicating a distinct microbial signature. Conversely, general HPV negativity corresponded to higher levels of classic oral commensals (*Haemophilus*, *Neisseria*), while oral HPV negativity was characterized by increased *Scardovia*, *Finegoldia*, *Shuttleworthia*, *Pseudomonas*, and *Propionibacterium*. Together, these patterns show that systemic HPV status and mucosal (oral) HPV detection correlate with different and non-overlapping oral microbiome signatures.

HDV seropositivity was significantly associated with increased abundance of *Aggregatibacter* (*Actinobacillus*), *Haemophilus*, *Veillonella*, *Fusobacterium*, and *Alloprevotella*. In contrast, HDV-negative individuals displayed a wider set of low-abundance environmental taxa, suggesting that in the absence of chronic hepatitis, oral communities may retain greater compositional diversity. Importantly, HDV-positive individuals exhibited significantly higher abundances of Aggregatibacter (*Actinobacillus*), of which many species of this genera are well-studied periodontal pathogens whose metabolic products induce strong pro-inflammatory cytokine responses such as IL-1β^73,74^, promoting periodontal tissue destruction^75,76^. Other enriched taxa in HDV-positive individuals included *Kingella*, *Haemophilus*, *Veillonella*, *Solobacterium*, *Eubacterium*, *Fusobacterium*, and *Alloprevotella*. HDV-negativitity associated genera included *Aliivibrio*, *Sphingomonas*, *Methylobacterium*, *Ralstonia*, *Brevundimonas*, many of which are environmental or transient members of the oral cavity.

For hepatitis B, active HBV infection showed correlation with higher levels of oral *Neisseria*, *Haemophilus*, *Alloprevotella*, and *Actinobacillus* (*Aggregatibacter*) compared to HBV-negative individuals (Fig. 6B, 6E). Given the negative associations with *Pelomonas*, *Xanthomonas*, *Propionibacterium*, and *Burkholderia* mirrored and smaller subset of the association patterns seen with HDV. One plausible explanation, although not testable here, is that HDV often co-occurs with HBV and behavioral or inflammatory risk factors that themselves correlate with reduced oral health^77^. The recurrent enrichment of periodontal-associated taxa in HDV/HBV-positive individuals supports a plausible link between viral hepatitis and their associated risk behaviors via the oral-liver axis^78^, and an oral microbiome shifted toward inflammatory anaerobes.

## Discussion

The human oral cavity is directly and repeatedly exposed to a broad spectrum of environmental stressors, including foods, aerosols, and medications. Prior work has identified strong associations and mechanistic links between oral communities and human health, spanning “local” outcomes such as gingivitis, periodontitis, and caries, as well as more distant associations with cardiometabolic and other systemic diseases^79,80^. To extend this literature, we leveraged public data from the National Health and Nutrition Examination Survey (NHANES) to build a resource for interrogating how the oral microbiome varies across multiple host phenotypes and exposures. By systematically querying associations between salivary microbiome composition and demographics, clinical variables, diseases, and a wide range of behavioral and chemical exposures, our study provides a population-scale view of how the oral microbiome in U.S. adults is structured by demography, life course, local oral conditions, the exposome, and systemic health.

A major implication of our findings is that demographic context and age are organizing axes of oral ecology. The stratification of taxa on coordinated non-linear age trajectories indicates that common oral may follow structured life-course dynamics, but this must be evaluated further. Our pseudo-temporal clustering of genera into age-“peaking” modules complements recent work showing that oral α-diversity peaks in early adulthood and that β-diversity increases across the lifespan.^17,81^ These patterns likely integrate changes in immunity, dentition, hormones, and accumulated environmental exposures.

Our analyses extend long-standing ecological models of periodontal disease and caries. The enrichment of well-characterized periodontal pathobionts—including *Porphyromonas*, *Prevotella*, *Tannerella*, and *Treponema*—in gum disease and poor oral hygiene is fully consistent with decades of subgingival plaque and periodontal literature.^27–31,37^ Likewise, the caries-associated network is enriched for acidogenic and saccharolytic genera such as *Bifidobacterium*/*Scardovia*/*Parascardovia*, *Lactobacillus*, *Actinomyces*, and *Olsenella*, mirroring culture-based and sequencing studies of root and enamel caries.^49,82–87^ At the same time, we identify less-recognized taxa such as *Eggerthia* and *Shuttleworthia*, that track with poor oral hygiene and periodontal status. They have been reported in association with gingivitis and periodontitis in large cohorts^88^. This suggests that large, well-powered population datasets can expand the cast of organisms that contribute to dysbiosis beyond the classic “red complex.” The distinct denture-associated niche we observe is in line with prior work showing that acrylic prostheses create a low-pH, plaque-like environment that favors caries-associated taxa and non-oral *Bifidobacteria.*^38–43,83^

Conversely, our identification of *Proteobacteria*-rich, oxygen-tolerant communities in caries-free and periodontally healthy individuals reinforces the notion that oral health is broadly characterized by aerobes and facultative anaerobes such as *Neisseria* and *Haemophilus*, previously recognized as markers of periodontal health and early colonizers in well-oxygenated plaque.^28,29,34–36^ The surprising negative association between caries and *Actinobacillus*/*Aggregatibacter* — a genus otherwise implicated in periodontal and invasive oral disease^32,33^ — highlights that some classically pathogenic genera may behave as low-abundance commensals in certain ecological contexts. This underscores the importance of distinguishing presence from ecological dominance, and suggests that the same taxon can occupy different functional roles depending on niche and community state.

Our exposome analyses show that the oral microbiome is potentially responsive to both behavioral and chemical exposures, with tobacco-related markers yielding the strongest and most coherent signals. Across serum biomarkers, urinary metabolites, and self-reported smoking, we observe a consistent difference from oxygen-tolerant commensals (e.g., *Neisseria*, *Haemophilus*, *Capnocytophaga*) toward aciduric and obligate anaerobes, including many genera implicated in caries and periodontitis. This pattern extends and clarifies earlier work showing smoking-associated depletion of *Proteobacteria* and enrichment of *Atopobium* and *Streptococcus* in U.S. adults.^17,21,44,89^ The graded changes we detect across smoking intensity and recency—where anaerobes accumulate with heavier exposure and recede within days of cessation—support a model in which tobacco rapidly alters local oxygen tension, pH, and inflammation, transiently favoring disease-associated communities. Similar but more heterogeneous signatures observed for halogenated solvents, PFAS, and other toxicants suggest that chronic chemical exposures may also perturb oral ecology; however, the low biomarker levels of these compounds make mechanistic interpretation more uncertain.

Dietary variation additionally was correlated to changes in oral communities. The strong positive correlations between sugar- and carbohydrate-rich intake and classic cariogenic genera—including *Lactobacillus*, *Bifidobacterium*, and *Scardovia*—reaffirm ecological plaque models in which saccharolytic metabolism and lactic-acid accumulation drive enamel demineralization and low-pH niche formation.^82–85^ Conversely, the broadly negative associations between higher dietary fiber and multiple disease-linked taxa—including *Parascardovia*, *Filifactor*, and *Mycoplasma*—are intriguing, as intervention studies have rarely reported such decreases in oral pathogens with fiber supplementation.^50,51^ Our finding that selenium intake and supplement use are associated with widespread suppression of *Firmicutes* and *Proteobacteria* is consistent with animal and gut microbiome data indicating that selenium modulates antioxidant and inflammatory pathways and interacts bidirectionally with microbial communities.^52–54^ It also echoes older reports linking selenium exposure to altered enamel properties and caries risk.^55–57^ Together, these patterns suggest that dietary factors can shift oral communities toward or away from dysbiotic states, though targeted interventional trials are needed to test causality and dose-response.

Finally, our systemic analyses indicate that oral microbial signatures extend beyond local pathology to correlations with respiratory, cardiovascular, inflammatory, and viral phenotypes. The shared enrichment of respiratory-associated genera such as *Pseudomonas* and *Acinetobacter* in asthma and emphysema is compatible with an oral–lung axis in which oropharyngeal communities contribute to airway colonization, inflammation, or both.^60–64,90^ Associations between leukocyte counts and periodontal/endodontic anaerobes such as *Desulfobulbus*, *Pseudoramibacter*, and *Cryptobacterium* align with prior work implicating these taxa in periodontitis, root-canal infection, and hyperglycemia-associated dysbiosis^66–70,72^, supporting the idea that elevated systemic inflammation may at least partly reflect increased oral anaerobic burden. The broad, distinct signatures associated with HPV, HBV, and HDV serologies—including recurrent enrichment of periodontal taxa in hepatitis virus–positive individuals—are consistent with emerging concepts of an oral–liver axis and shared behavioral and inflammatory pathways linking viral hepatitis, oral dysbiosis, and liver disease.^77,78^

The majority of microbiome association studies have focused on the lower gastrointestinal tract; by comparison, the oral microbiome has been markedly understudied, particularly in diverse populations^12^. Our results argue that the oral microbiota warrants similar attention, both for its potential contributions to disease and for its utility as a minimally invasive indicator of health or even therapeutic target. Prior work has estimated the variance in host phenotypes explained by gut microbial composition; applying analogous variance-partitioning approaches to oral communities, particularly in designs that directly compare oral versus gut-associated phenotypes^91^, could clarify the relative information content of these ecosystems and help prioritize future diagnostic development. At a minimum, larger and more deeply profiled cohorts using modern sequencing technologies (e.g., shotgun metagenomics rather than 16S alone) are needed to resolve functional variation in oral communities and to determine how such functional shifts associate with, and potentially drive, health and disease.

In summary, we believe our results highlight the promise of the oral cavity as both a window into, and a potential lever for, diseases that are local to the mouth as well as systemic. With future, larger-scale efforts that integrate more diverse populations, deeper functional profiling, and comparative analyses across body sites, we expect links between the oral microbiome, environmental exposures, disease, and host health to be further solidified and increasingly translated into real-world impact.

### Limitations of the study

The cross-sectional, observational nature of NHANES precludes causal inference and cannot distinguish whether microbiome changes are drivers, consequences, or correlates of host phenotypes. Although we adjusted for multiple comparisons (controlling the false discovery rate), the complexity of the sample population likely means that latent confounders remain and that a subset of associations may represent false positives.

Oral profiling was based on single 16S rRNA amplicon samples, limiting taxonomic resolution to the genus level and obscuring strain-level heterogeneity in virulence and metabolism. In addition, NHANES profiled saliva, which aggregates planktonic cells from multiple oral micro-ecosystems and is sensitive to short-term behaviors (e.g., time since last toothbrushing). The oral microbiome is known to exhibit strong spatial heterogeneity across niches such as dental plaque, tongue dorsum, and mucosal surfaces, and our salivary measurements do not capture this fine-scale ecological structure.

Key exposures, particularly diet, were assessed from single 24-hour recalls, introducing potential misclassification and reducing sensitivity to long-term patterns. Some associations may also be sensitive to the choice of compositional data transformation; for example, we observed heterogeneity in effect estimates between relative-abundance and centered log-ratio–transformed data, although the strongest findings were conserved. Despite extensive adjustment for demographic and behavioral covariates and the use of survey weights, residual confounding from unmeasured factors—such as detailed oral hygiene practices, recent dental interventions, and medication use—is likely.

Finally, our findings are anchored to the U.S. population context and may not generalize to settings with different environmental, cultural, or healthcare landscape, and tissue samples from other parts of the human host, such as the lung, skin, or gut; more globally diverse cohorts will be essential to establish external validity. Addressing these gaps will require longitudinal designs, sampling the same person across tissue type, shotgun metagenomics and metabolomics for functional resolution, comparisons across oral sampling sites, and integration with gut microbiome and host genomic data to clarify causal pathways.

## Supplementary Figures

**Figure S1.**
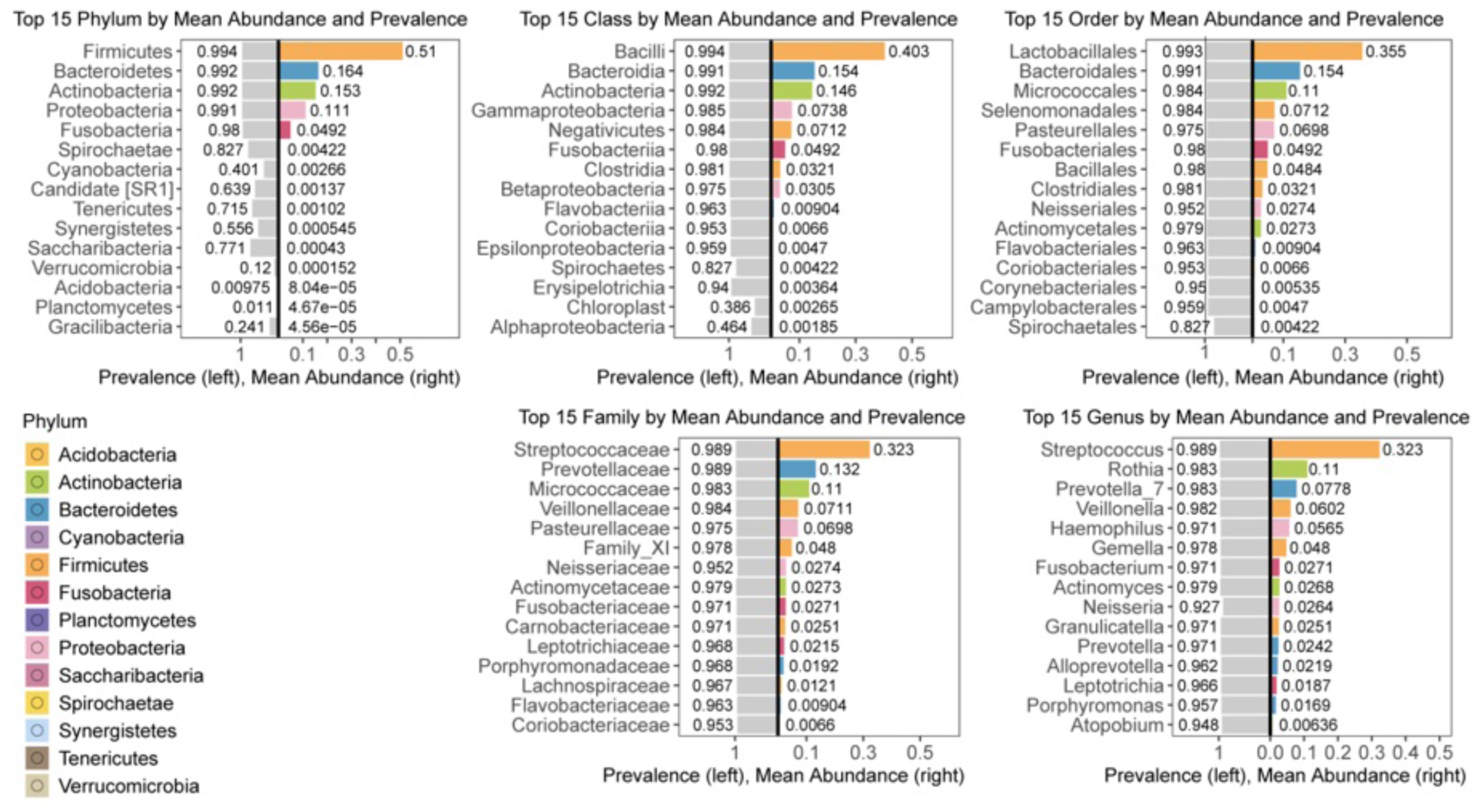
Taxonomic Composition of the Oral Microbiome. Mean relative abundance and prevalence of the top 15 taxa at each rank (phylum, class, order, family, and genus). Right panels show mean relative abundance; left panels show prevalence across participants. Phylum-level color coding is consistent across all plots.

**Figure S2.**
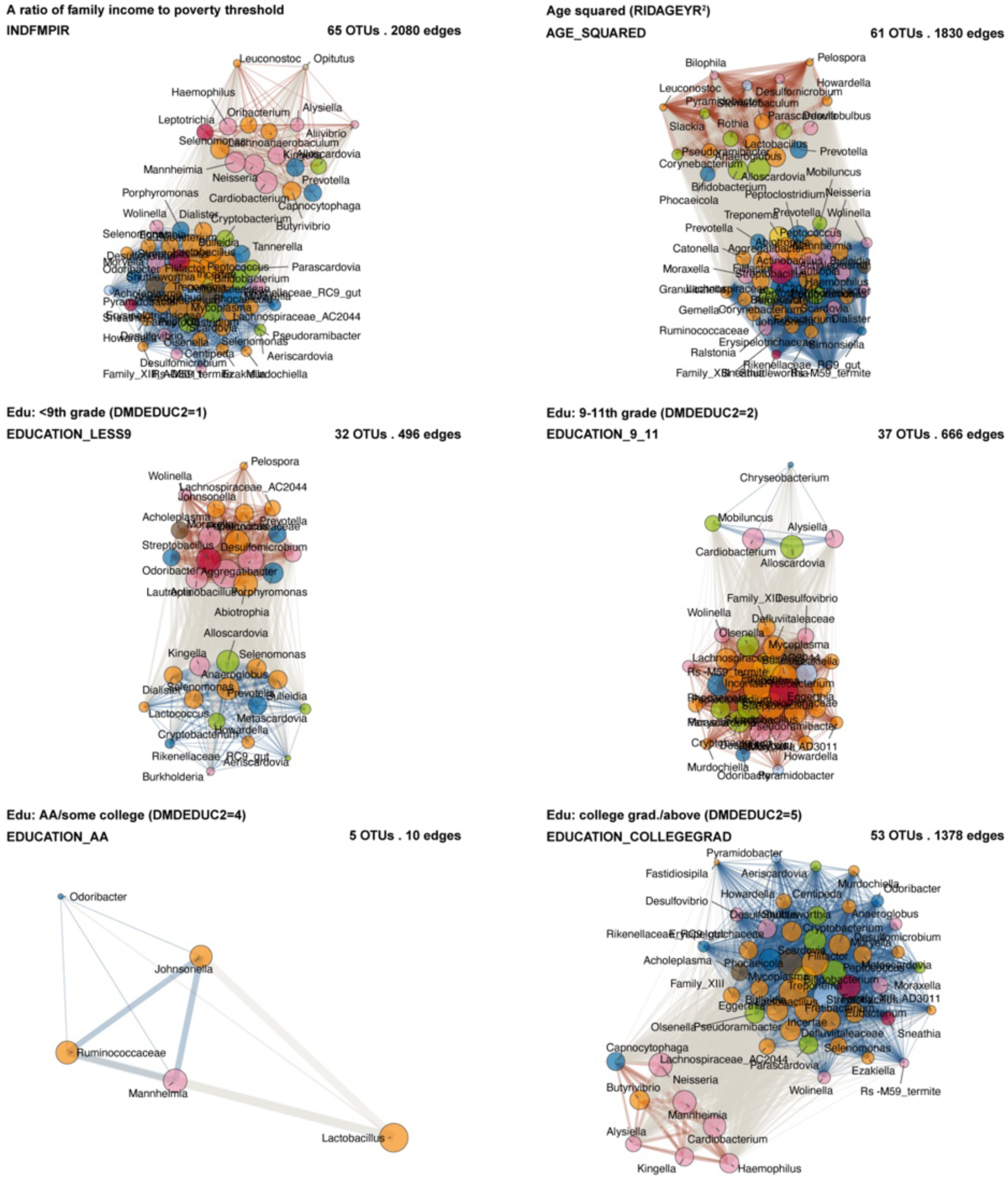
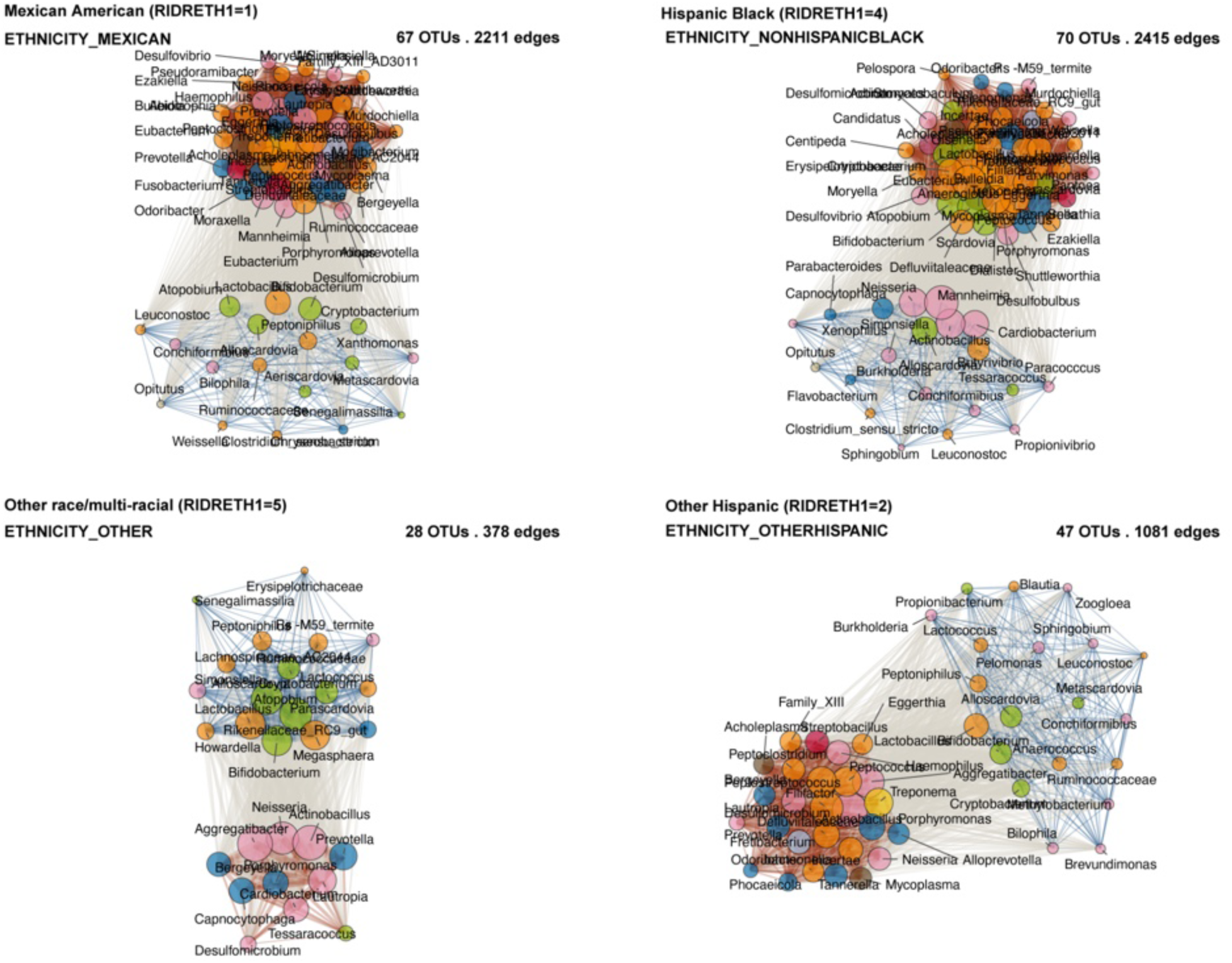
Full list of association effect networks of CLR-transformed common oral genera with demographic variables. Survey-weighted linear models were fitted between CLR-transformed oral genera and demographic variables across two NHANES cycles (2009–2010, 2011–2012). Networks include only variables with at least two significant genera passing FDR ≤ 0.05, q ≤ 0.05, and the 10% prevalence threshold. Node size represents the cumulative absolute effect size (|β|), and node color indicates phylum. Edges link genera with concordant associations; width reflects the geometric mean of effect sizes, and color encodes direction (blue = negative, orange = positive).

**Figure S3.**
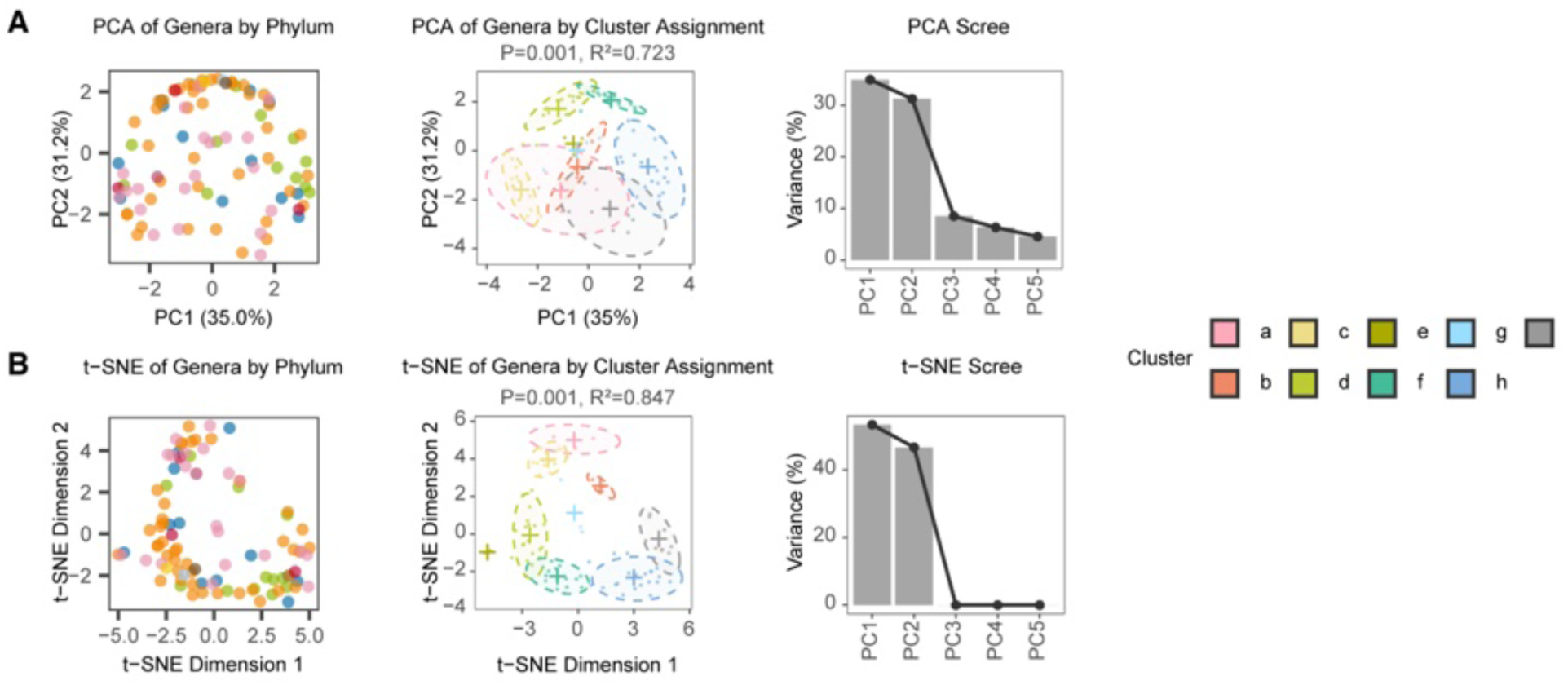
Principal component and t-SNE result of age-associated oral-community clusters. (A) Principal component analysis (PCA) of 93 prevalent genera. Left: PCA of genera colored by cluster assignment, with PC1 and PC2 axes labeled with variance explained. Middle: PCA of genera colored by cluster assignment, with PERMANOVA p value and R² for cluster separation indicated. Right: scree plot showing variance explained by the first five principal components. (B) t-SNE embedding of the same genera, with three corresponding panels (left, middle, right) mirroring the PCA layouts in (A): clusters colored on the t-SNE map, cluster-level separation statistics (p value and R²), and variance explained by the first five t-SNE dimensions.

**Figure S4.**
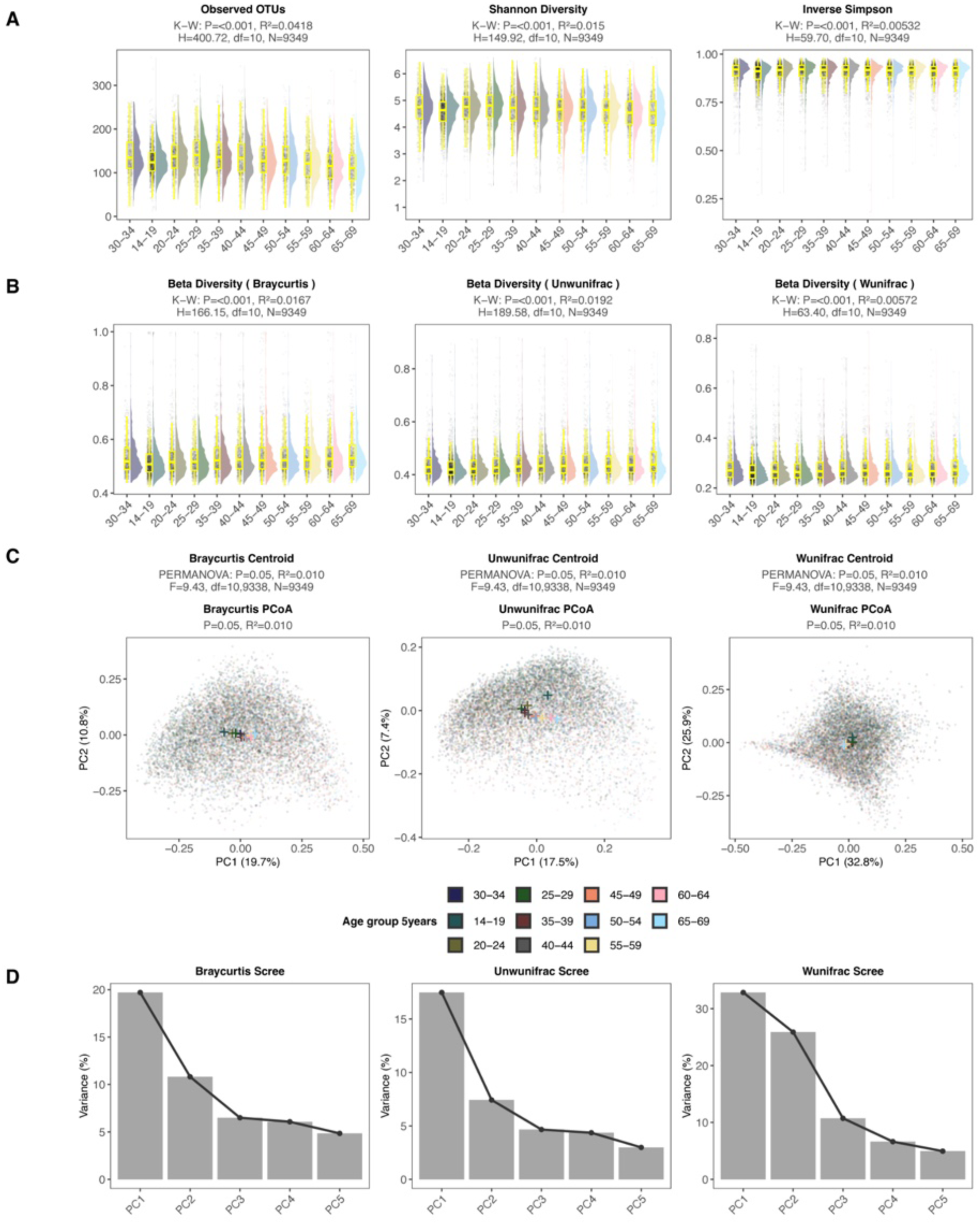
Age-stratified α- and β-diversity and distribution. Summary statistics are shown; full pairwise comparisons are provided in Supplementary Tables S5, S6, and S7. (A) Box-and-violin plots of α-diversity (Observed OTUs, Shannon index, inverse Simpson index; left to right) across 5-year age groups (14–19 to 65–69 years; reference group leftmost). (B) Box-and-violin plots of β-diversity (Bray–Curtis, unweighted UniFrac, weighted UniFrac; left to right) across the same age groups (reference group leftmost). (C) Principal coordinate plots for Bray–Curtis, unweighted UniFrac, and weighted UniFrac (left to right), with points colored by 5-year age group. (D) Scree plots of variance explained by the first five principal coordinates for Bray–Curtis (left), unweighted UniFrac (middle), and weighted UniFrac (right).

**Figure S5.**
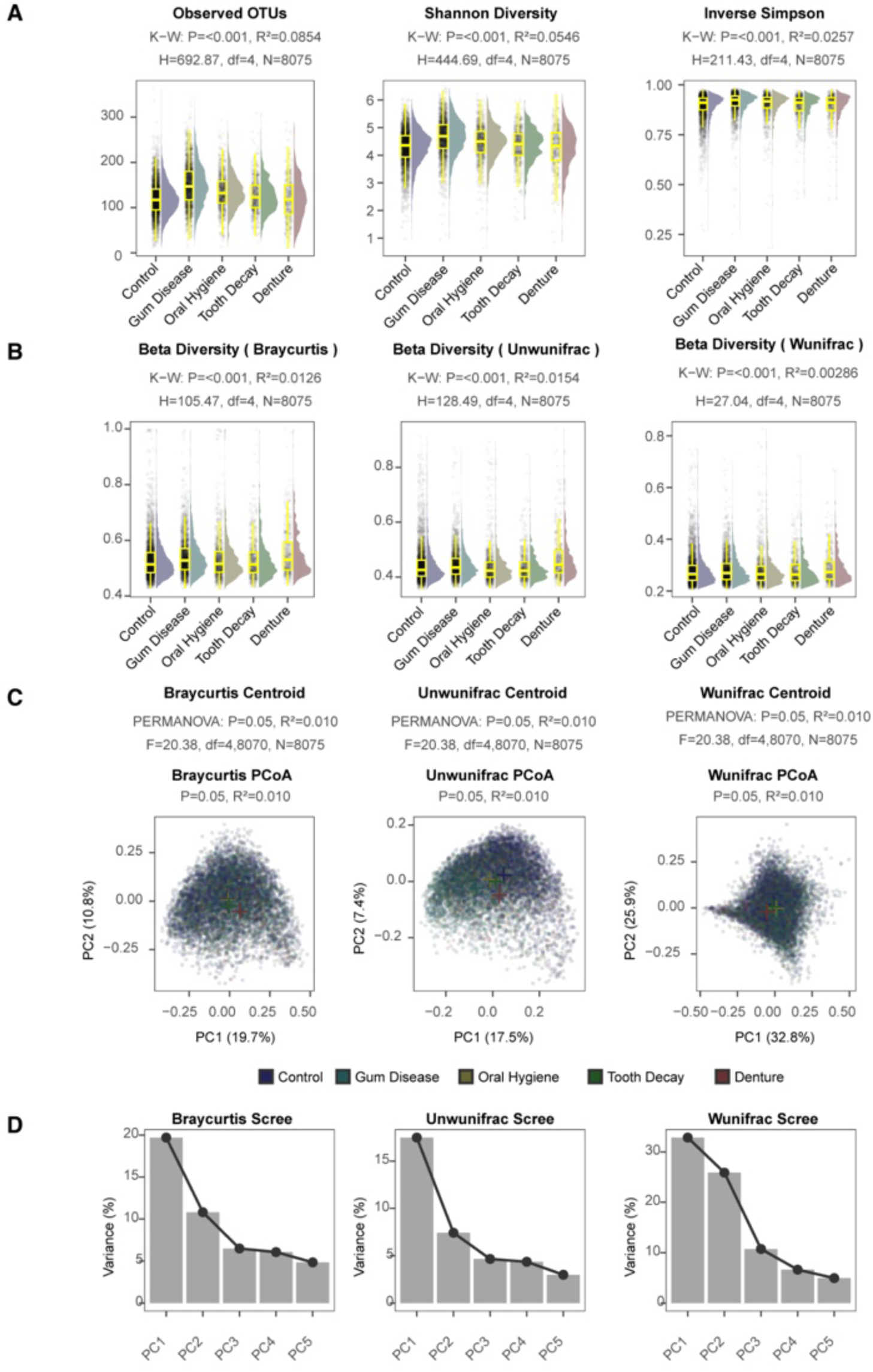
Local oral disease group stratified α- and β-diversity distribution. Summary statistics are shown; full pairwise comparisons are provided in Tables S8, S9, and S10. (A) Box-and-violin plots of α-diversity (Observed OTUs, Shannon index, inverse Simpson index; left to right) across oral disease categories (control [leftmost], gum disease, oral hygiene needed, tooth decay, denture). (B) Box-and-violin plots of β-diversity (Bray–Curtis, unweighted UniFrac, weighted UniFrac; left to right) across the same oral disease categories (control leftmost). (C) Principal coordinate plots for Bray–Curtis, unweighted UniFrac, and weighted UniFrac (left to right), with points colored by oral disease category. (D) Scree plots of variance explained by the first five principal coordinates for Bray–Curtis (left), unweighted UniFrac (middle), and weighted UniFrac (right).

**Figure S6.**
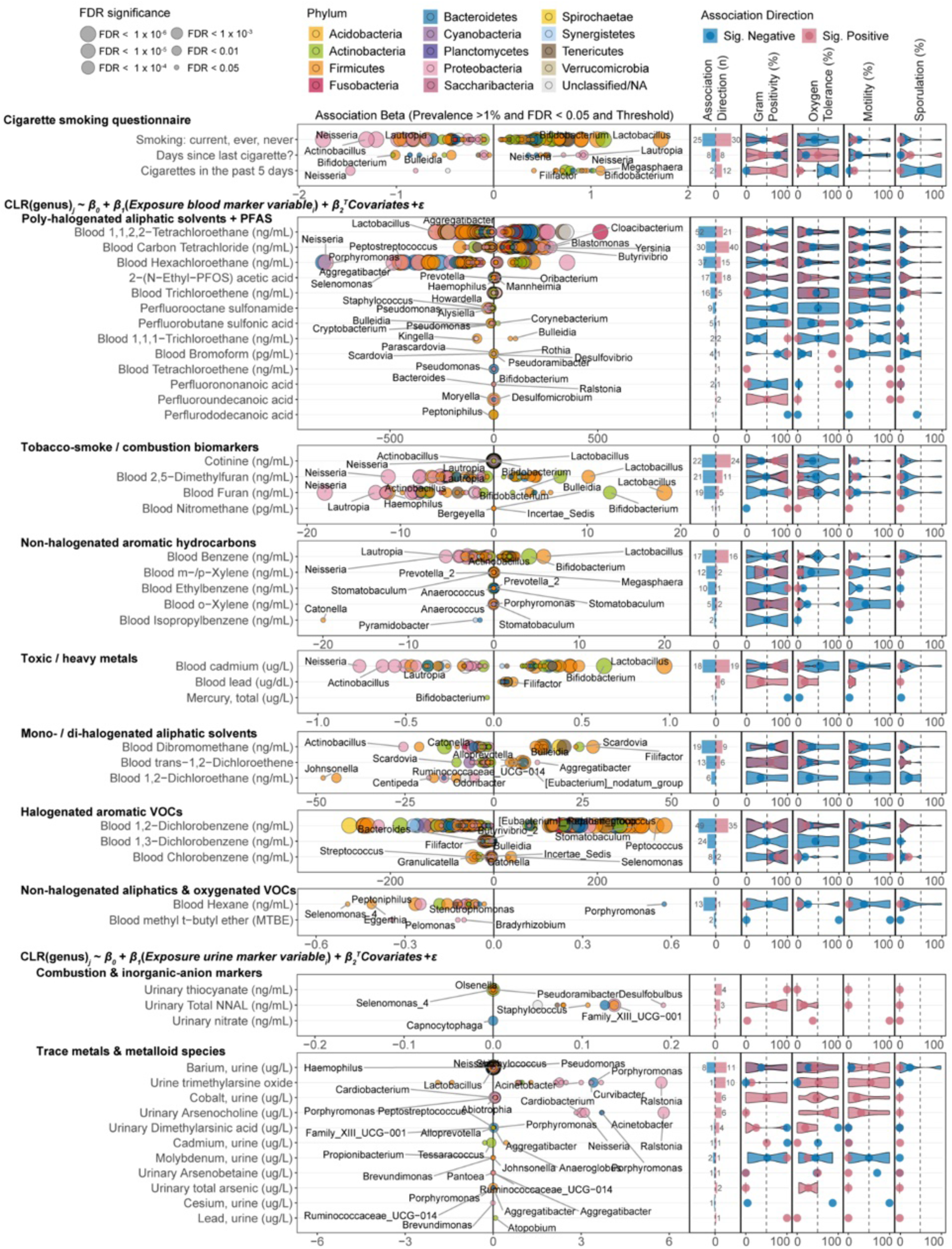
Associations of oral genera with smoking-related questionnaire, blood, and urinary markers. Survey-weighted regressions across two NHANES oral microbiome cycles (2009–2010, 2011–2012) relating CLR-transformed genera to smoking and combustion-related questionnaire items and biomarker concentrations. Left (“Terry plots”): regression coefficients β₁ from models of the form CLR(genus)ⱼ ∼ β₀ + β₁(Exposure markerᵢ) + β₂ᵀCovariates + ε, for blood or urinary exposure variables (prevalence ≥1%, FDR ≤0.05, q ≤0.05), with dot size indicating FDR category. Right: GOLD-DB–derived microbial phenotype indices (Gram positivity, oxygen tolerance, motility, sporulation) summarized by association direction (positive/negative). Only exposure variables with at least one significant, GOLD-DB–annotated genus are shown.

**Figure S7.**
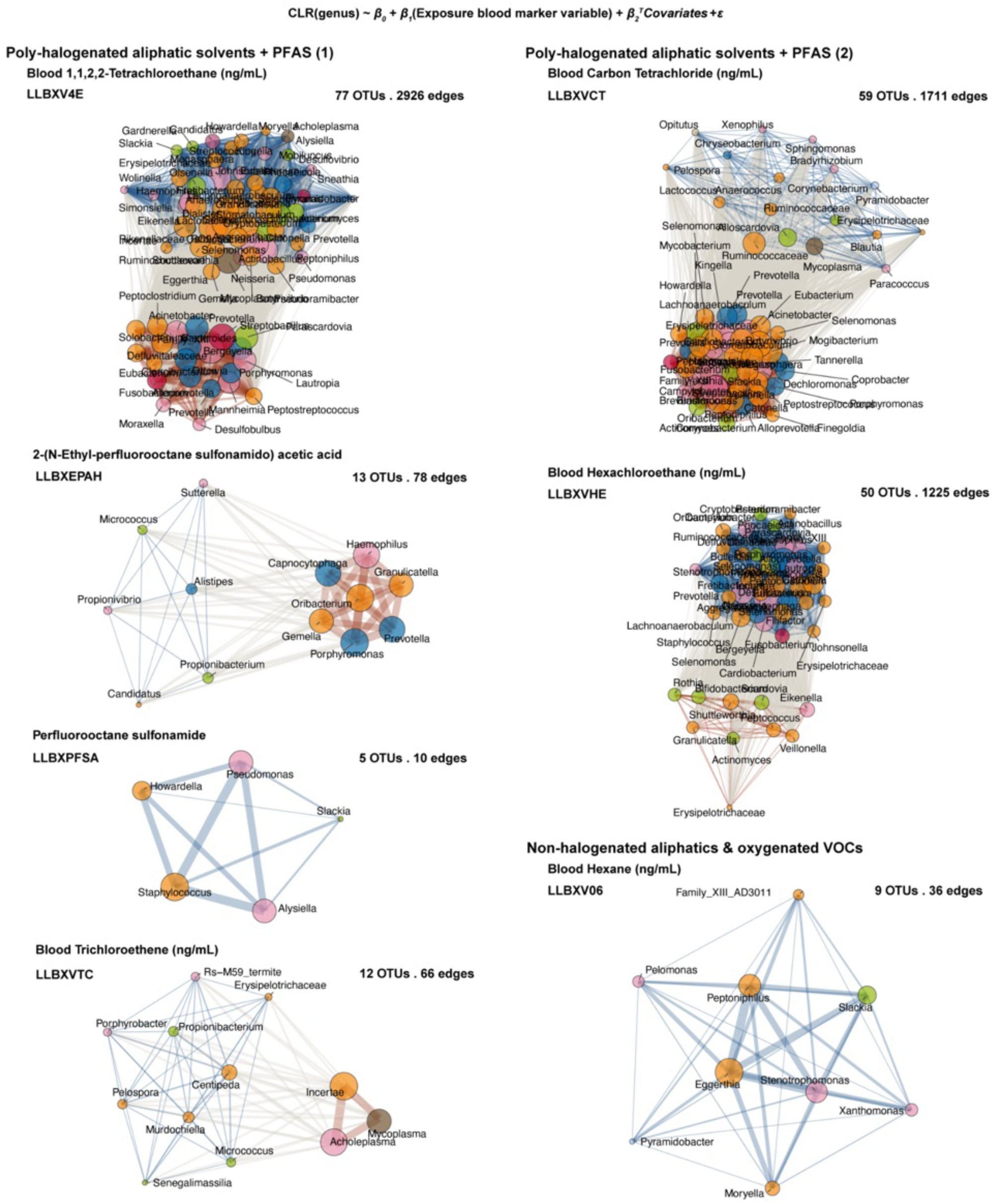
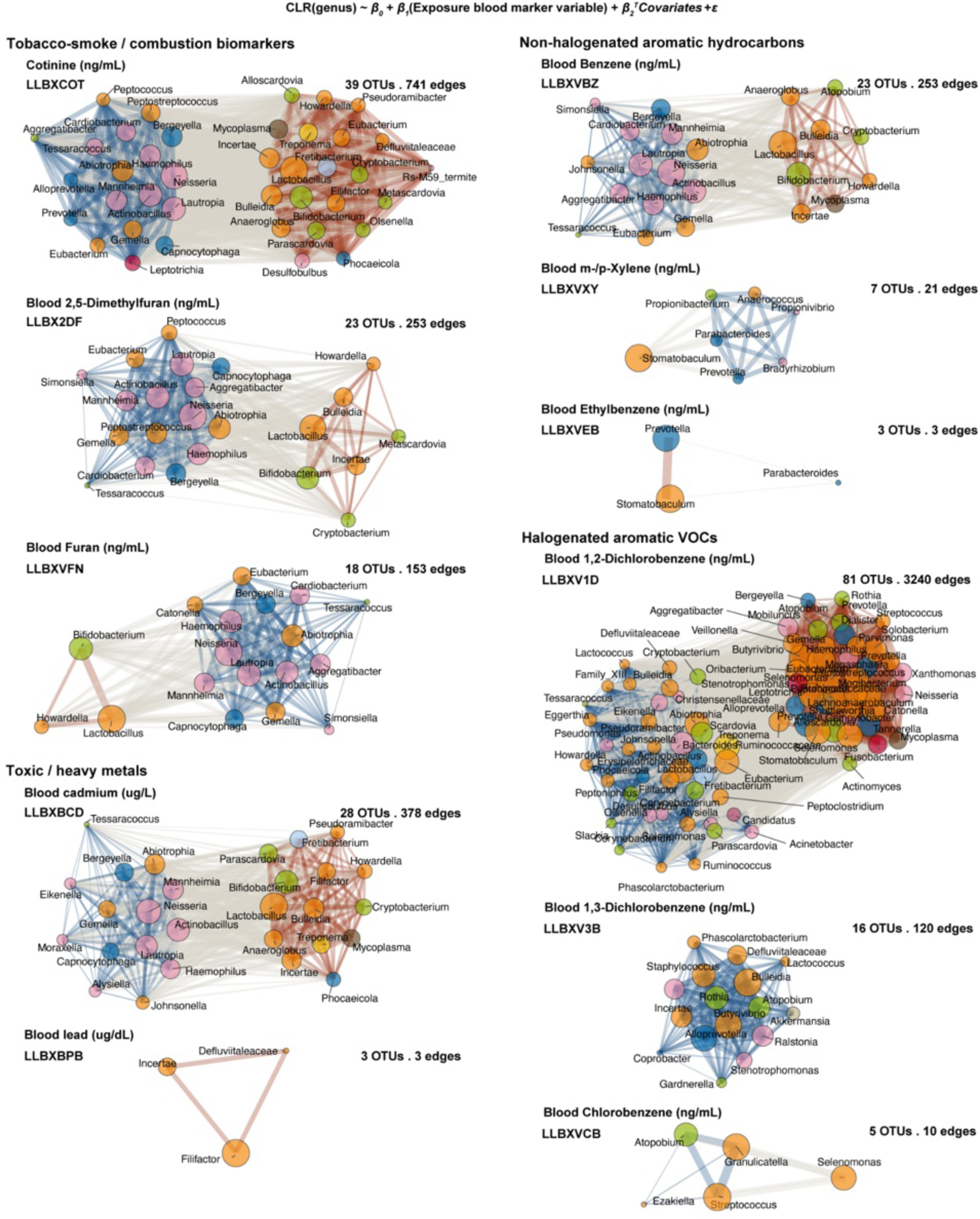
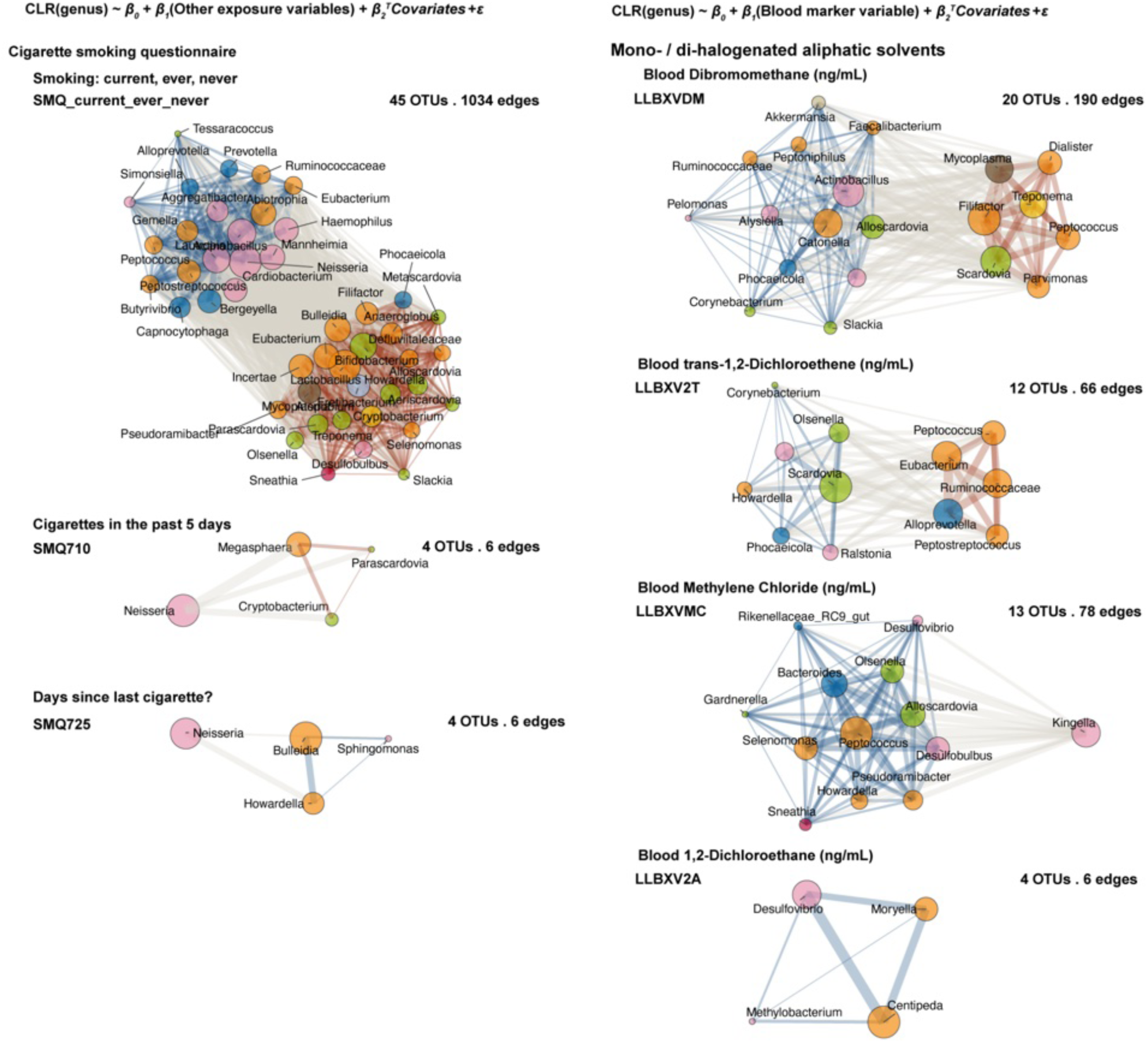
Networks of oral genera associated with combustion-related exposures. Survey-weighted linear models related CLR-transformed genera to combustion-related exposure variables across NHANES 2009–2010 and 2011–2012: CLR(genus)ⱼ ∼ β₀ + β₁(Exposureᵢ) + β₂ᵀCovariates + ε. Networks include only exposures with ≥2 significant genera (prevalence ≥10%, FDR ≤0.05, q ≤0.05). Node size reflects cumulative |β|, node color indicates phylum, and edges connect genera with concordant effects; edge width is the geometric mean of effect sizes and edge color denotes direction (blue, negative; orange, positive). Figure S7 (continued) displays additional exposure-specific networks.

**Figure S8.**
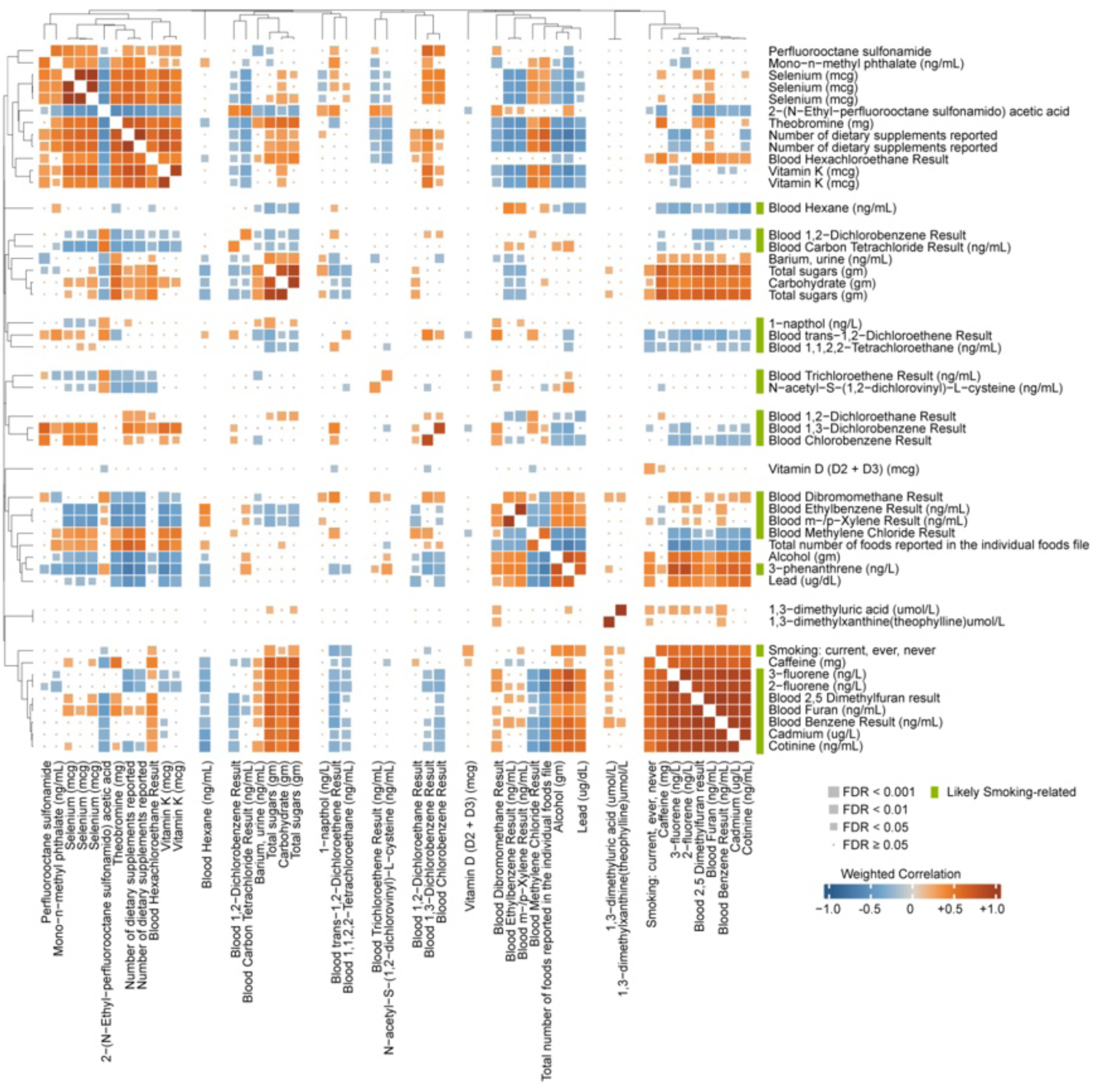
Smoking and combustion related exposure signatures in the oral microbiome. Weighted correlation matrix of microbial association signatures across smoking/combustion variables, restricted to exposures with ≥5 shared significant OTUs (prevalence ≥10%, FDR ≤0.01). Square size indicates FDR category and color indicates correlation strength. Hierarchical clustering (Euclidean distance) defines nine exposure clusters; right-side row annotation marks variables labeled as “Likely smoking-related” (light green).

**Figure S9.**
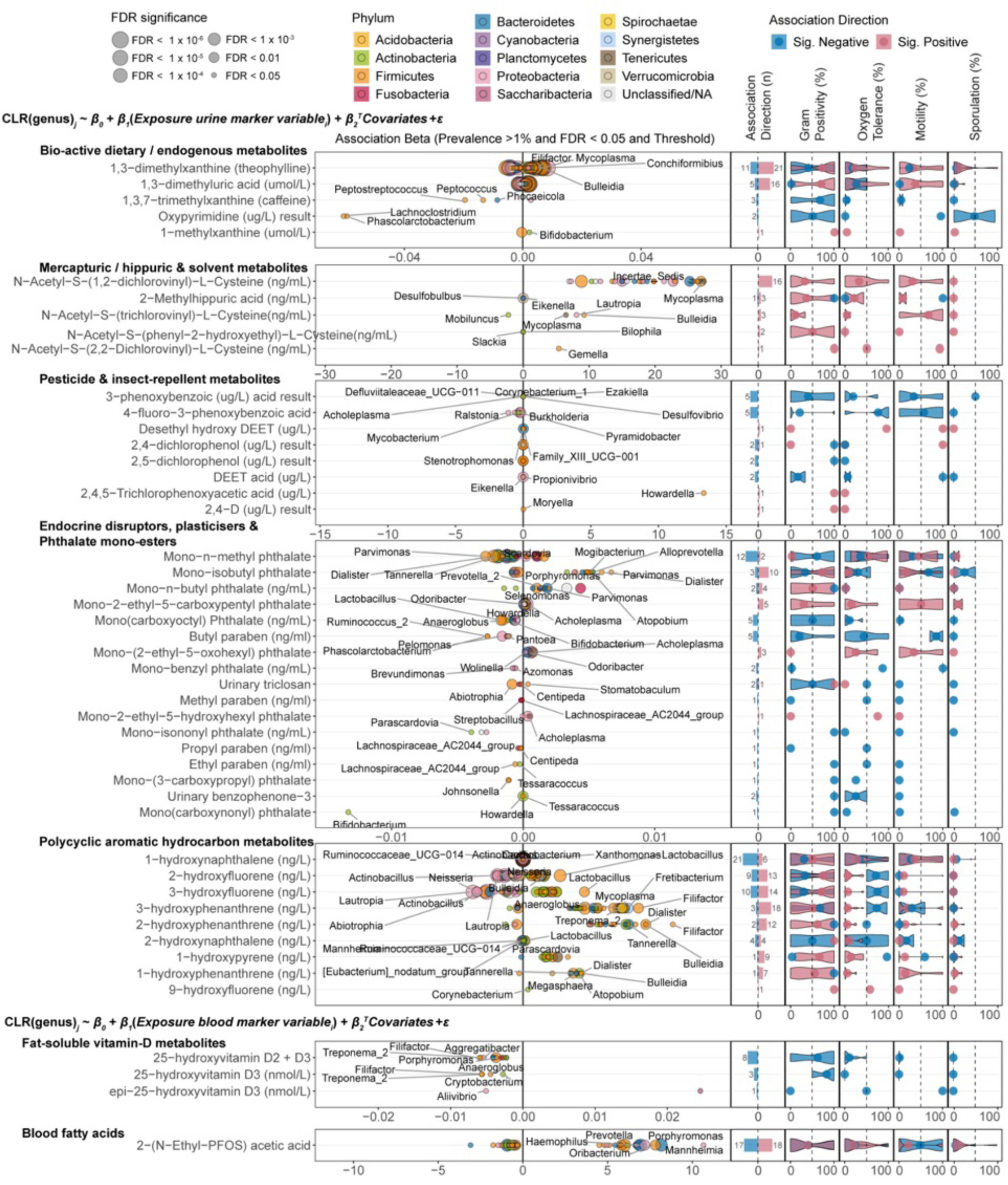
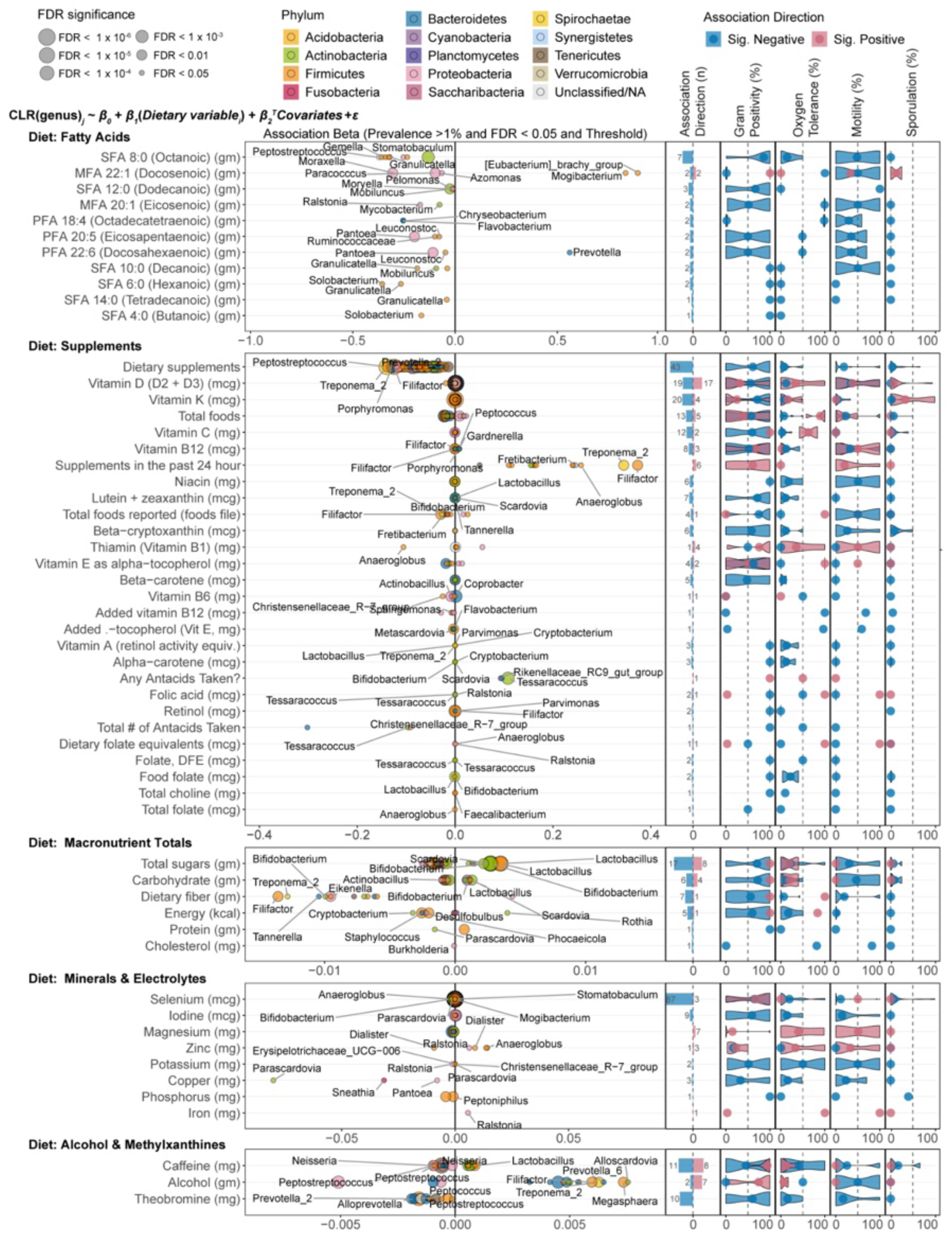
Oral genera associated with non–smoking-related urinary, blood, and dietary exposure markers. Survey-weighted regressions (NHANES 2009–2010, 2011–2012) relating CLR-transformed genera to urinary, blood, and dietary exposures: CLR(genus)ⱼ ∼ β₀ + β₁(Exposureᵢ) + β₂ᵀCovariates + ε. Left (“Terry plots”): β₁ coefficients for significant associations (prevalence ≥1%, FDR ≤0.05, q ≤0.05), with dot size indicating FDR category. Right: GOLD-DB microbial phenotype indices (Gram positivity, oxygen tolerance, motility, sporulation) summarized by association direction; only exposures with at least one significant GOLD-DB–annotated genus are shown.

**Figure S10.**
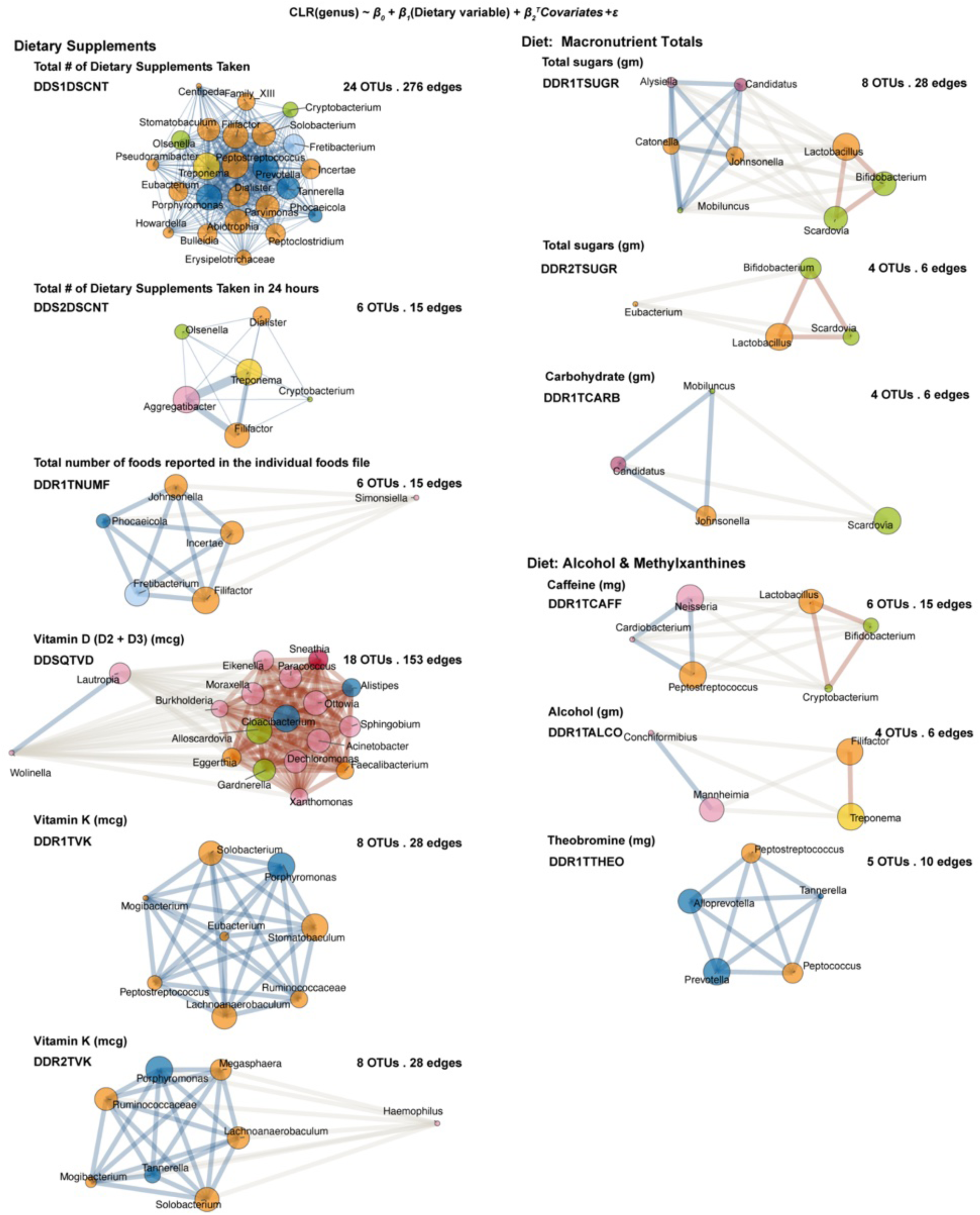
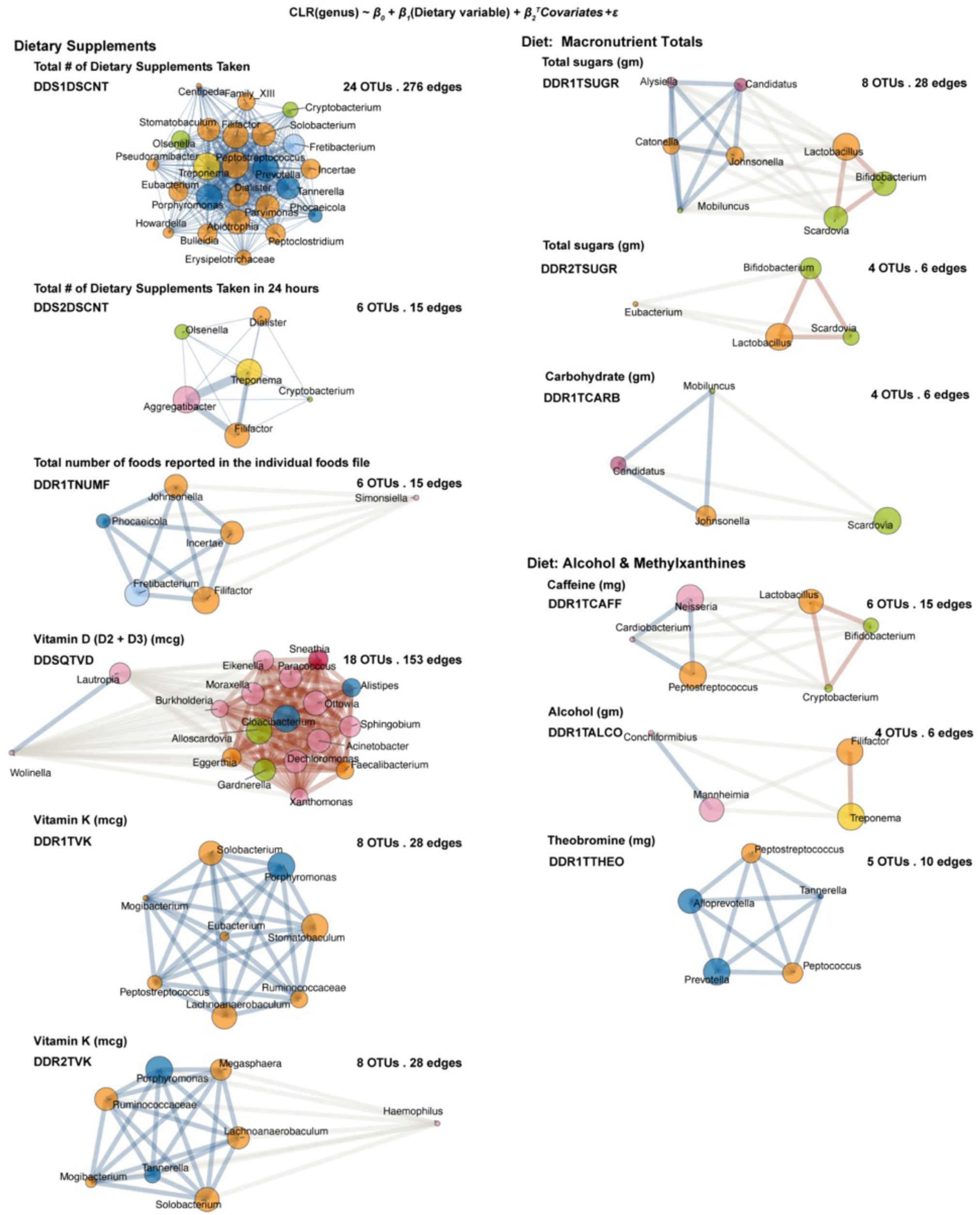
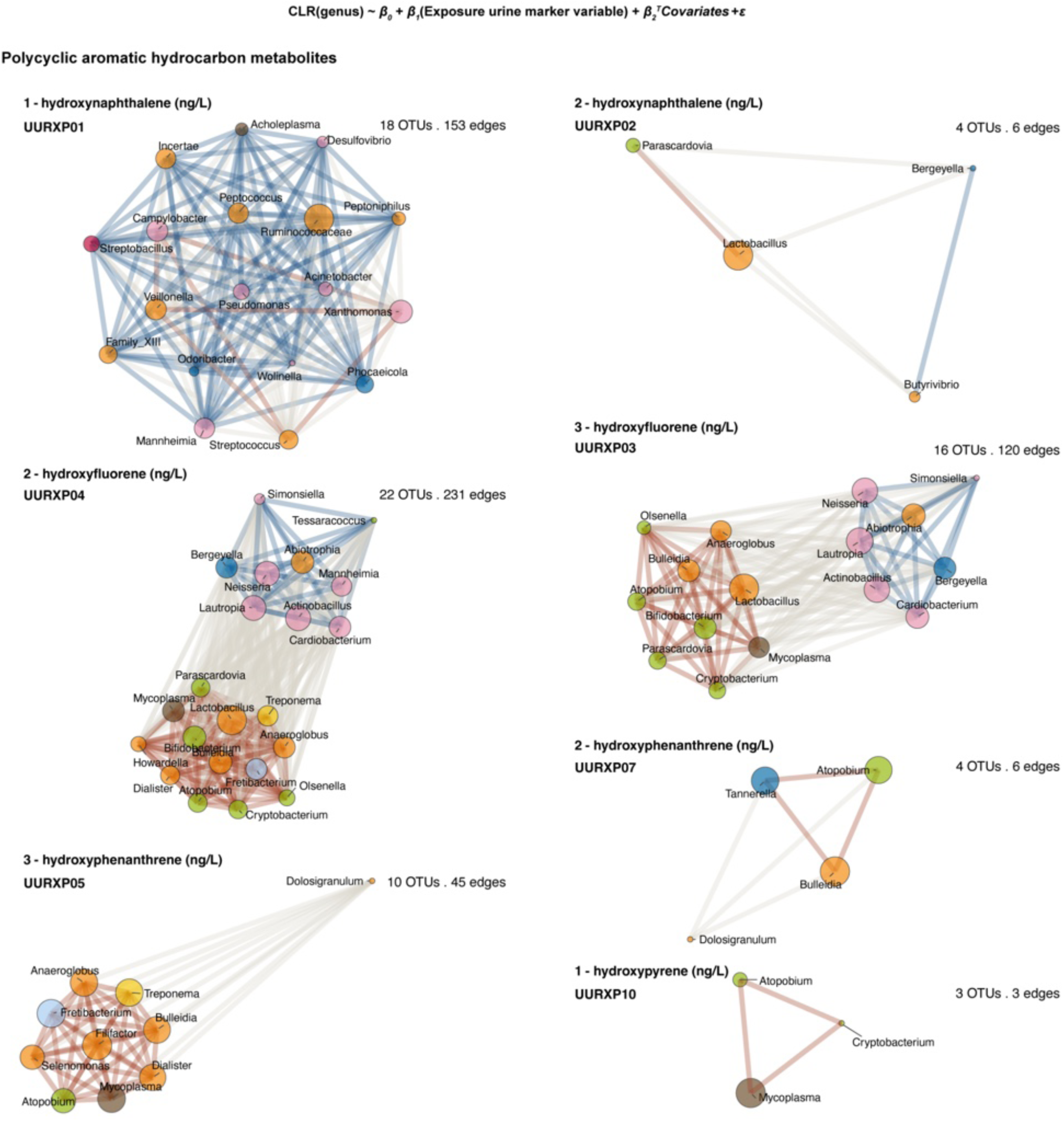
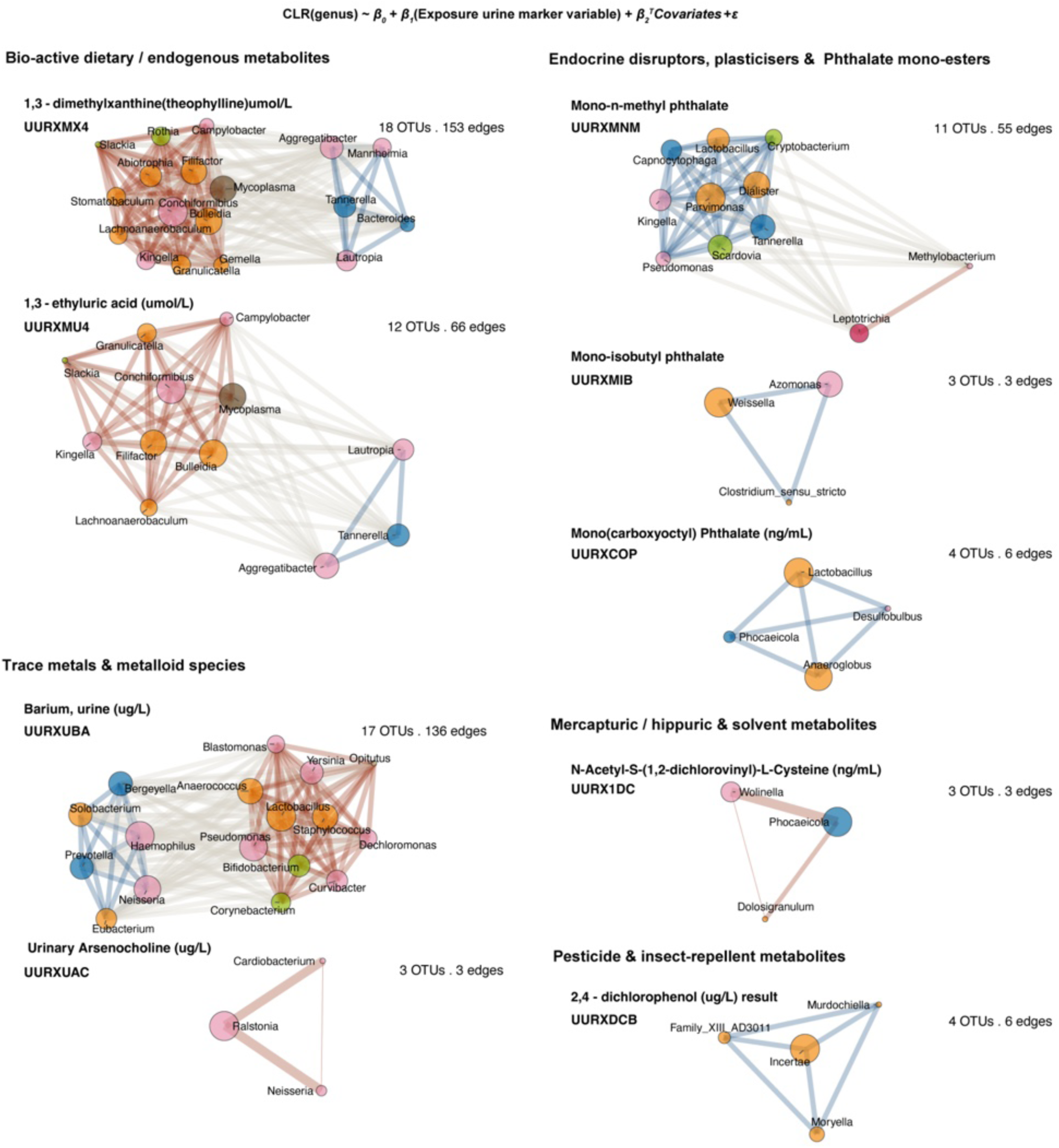
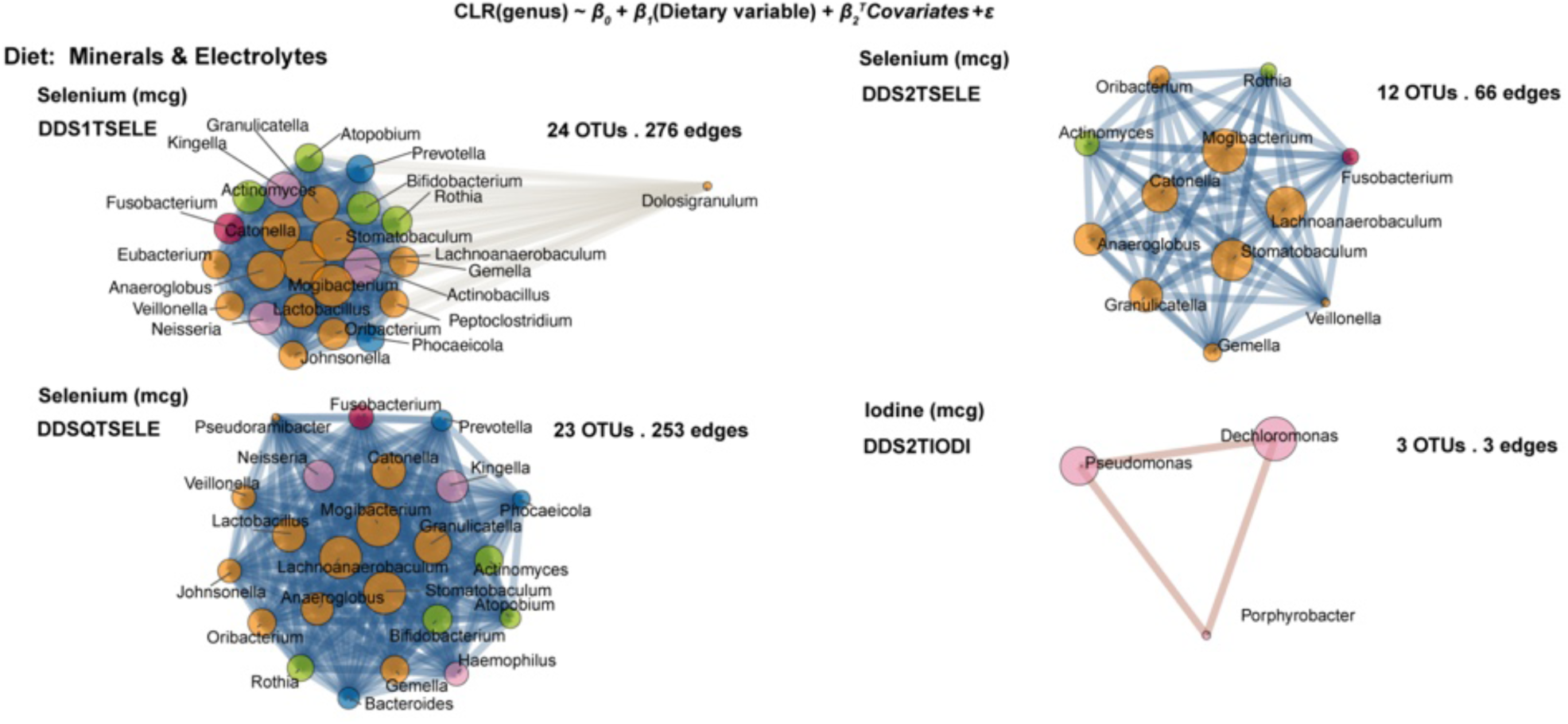
Small-effect association networks of common oral genera with non–smoking-related urinary, blood, and dietary exposures. Survey-weighted linear models related CLR-transformed genera to dietary exposure variables (prevalence ≥10%, FDR ≤0.05, q ≤0.05). Networks are shown for all dietary markers meeting these criteria. Nodes represent genera, with size proportional to cumulative absolute effect size (|β|) and color indicating phylum. Edges connect genera with concordant associations; edge width reflects the geometric mean of effect sizes, and edge color denotes direction (blue, negative; orange, positive).

**Figure S11.**
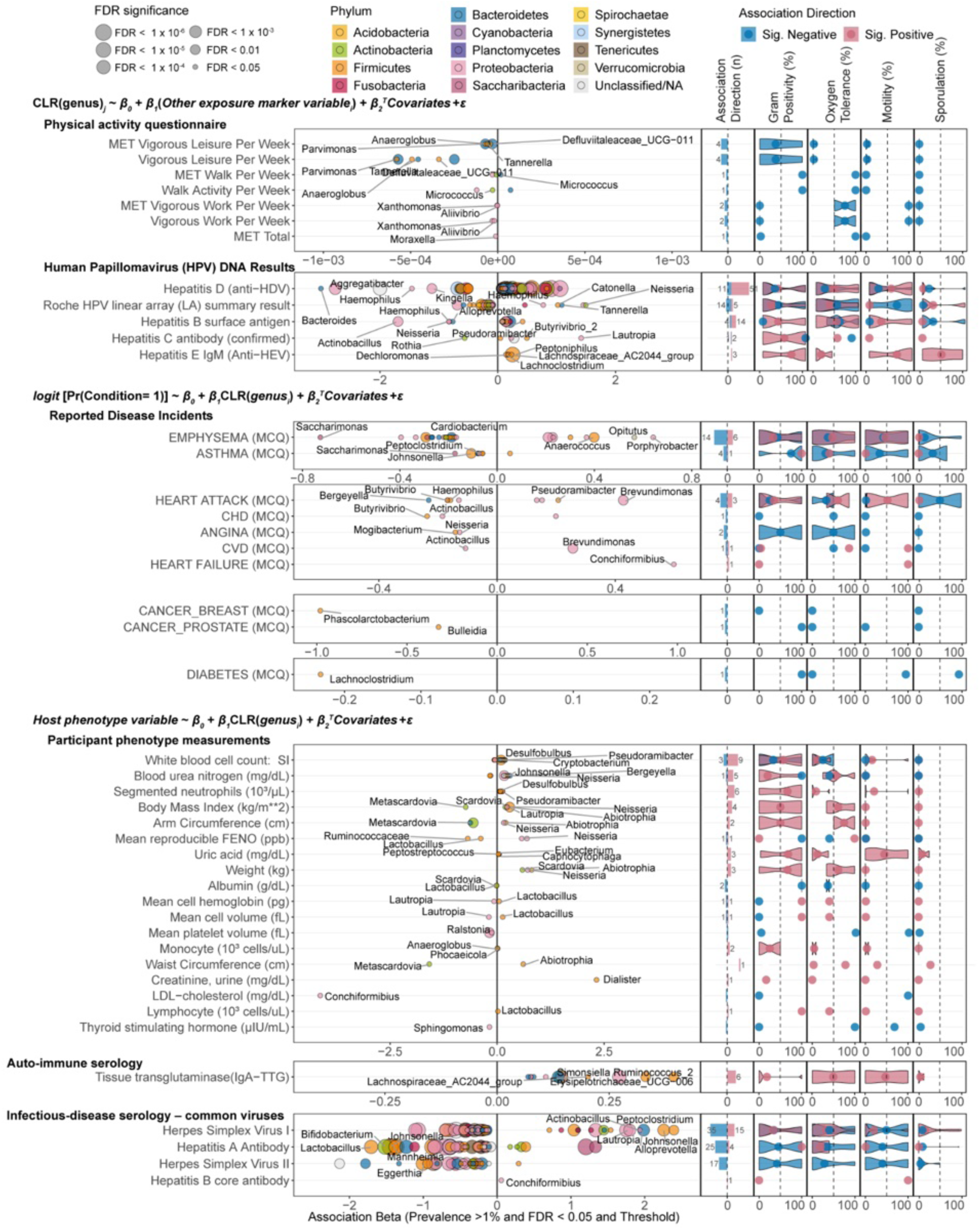
Significant associations of CLR-transformed oral genera with host exposures, diseases, and phenotypes. Survey-weighted regressions across two NHANES oral microbiome cycles (2009–2010, 2011–2012). (A) Associations with other exposure markers, from models CLR(genus)ⱼ ∼ β₀ + β₁(Other exposure variableᵢ) + β₂ᵀCovariates + ε. (B) Associations with reported disease incidents, from logistic models logit[Pr(Condition = 1)] ∼ β₀ + β₁·CLR(genusᵢ) + β₂ᵀCovariates + ε. (C) Associations with participant phenotype measurements, from models Host phenotype variable ∼ β₀ + β₁·CLR(genusᵢ) + β₂ᵀCovariates + ε. Left (“Terry plots”): regression coefficients β₁ for significant associations (prevalence ≥1%, FDR ≤0.05, q ≤0.05), with dot size indicating FDR category. Right: GOLD-DB–derived microbial phenotype indices (Gram positivity, oxygen tolerance, motility, sporulation) summarized by association direction (positive/negative). Only variables containing at least one significant GOLD-DB–annotated genus are shown.

**Figure S12.**
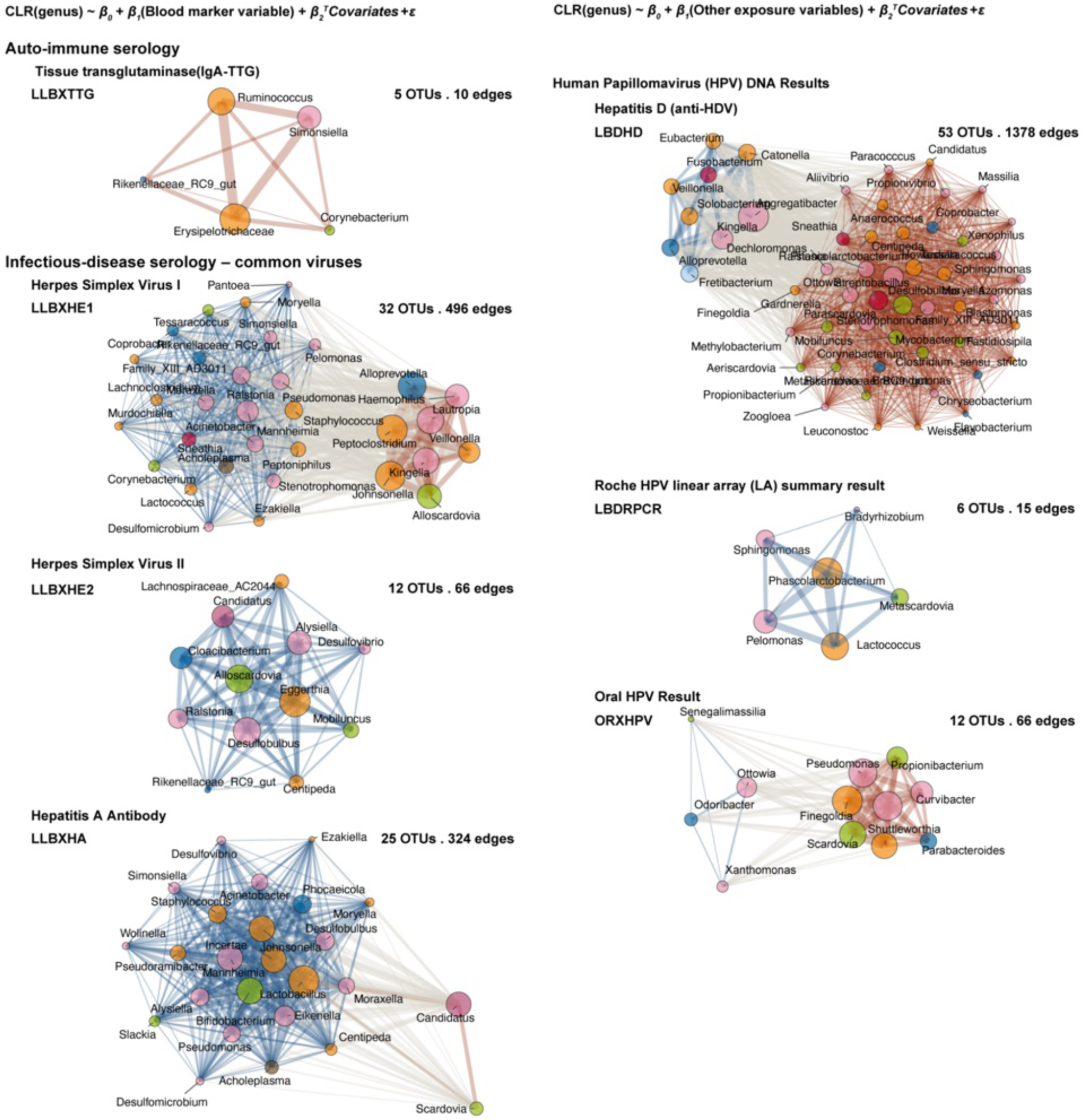
Extended list of small-effect networks for host exposures, diseases, and phenotypes. Network representations of significant associations between CLR-transformed common oral genera and host exposures, reported diseases, and phenotype measurements (prevalence ≥10%, FDR ≤0.05, q ≤0.05). Nodes represent genera, with size proportional to cumulative absolute effect size (|β|) and color indicating phylum. Edges connect genera with concordant associations; edge width reflects the geometric mean of effect sizes, and edge color denotes direction (blue, negative; orange, positive).

## Supplementary Tables

**Table S1 Full list of NHANES host variables, definitions, and distributions.** Comprehensive listing of all demographic, behavioral, dietary, exposure, clinical, and disease variables included in the Microbiome Association Study (MAS).

**Table S2 Centered log-ratio (CLR) microbiome-wide association results.** Complete survey-weighted regression results for associations between CLR-transformed genus-level abundances and host variables across all five modeling schemas described in Table 1. The table includes host variable, microbial taxon, regression coefficient (β), standard error, p value, Benjamini–Hochberg FDR, Storey q value, indicator for taxa meeting prevalence threshold when applicable, and model specification. Results correspond to analyses described in Table 1 and STAR Methods.

**Table S3 Relative abundance microbiome-wide association results.** Full survey-weighted association results using non-transformed relative abundance data (as produced by NHANES). Table structure mirrors Table S2, including regression coefficients, standard errors, p values, FDR, q values, and model identifiers.

**Table S4 GOLD database mapping and genus-level phenotype index aggregation.** Integration of NHANES oral genera with microbial phenotype annotations from the Genomes OnLine Database (GOLD v10). Table reports original and harmonized genus names, mapping status, number of matched species, intra-genus trait agreement level (high/medium/low), and aggregated phenotype indices (Gram stain, motility, sporulation, oxygen requirement).

**Table S5 Alpha- and beta-diversity distributions across defined host groups.** Survey-weighted distributions of alpha diversity (observed richness, Faith’s PD, Shannon, Simpson) and beta diversity (Bray–Curtis, weighted and unweighted UniFrac) stratified by demographic, exposure, and disease categories. Includes weighted means, dispersion metrics, and global test statistics.

**Table S6 Pairwise Wilcoxon tests for beta-diversity comparisons.** Complete pairwise group comparisons for beta-diversity metrics. Table reports test statistic, raw p value, and Benjamini–Hochberg–adjusted FDR for each group contrast.

**Table S7 Pairwise Wilcoxon tests for alpha-diversity comparisons.** Complete pairwise comparisons for alpha-diversity metrics across defined host categories. Includes test statistic, raw p value, and Benjamini–Hochberg–adjusted FDR-adjusted p value for each comparison.

**Table S8 Age cluster determination diagnostics.** Full results for data-driven clustering of age-associated genera. Includes silhouette widths across candidate k values, gap statistic estimates with bootstrap reference distributions, cluster stability metrics, and PERMANOVA statistics validating cluster separation.

**Table S9 Age cluster assignments for prevalent oral taxa.** Cluster membership assignments for taxa detected in ≥10% of participants and included in age trajectory analysis. Table reports genus name, phylum, standardized abundance trajectories across age bins, cluster identity (k = 9), and summary trajectory characteristics.

## Lead contact

Further information and requests for resources and reagents should be directed to and will be fulfilled by the contacts, Byeongyeon (Terry) Cho, Braden Tierney, and Chirag Patel.

## Materials availability

This study did not generate new or unique biological reagents. All raw and processed data used herein were derived from publicly available sources generated and released by Vogtmann et al. (2023) and the National Center for Health Statistics^15,16,92,93^. The fully processed dataset, including taxonomic abundances and integrated microbial phenotype annotations from GOLD-DB, generated as part of this study, is deposited and publicly available at https://doi.org/10.5281/zenodo.17849473.

## Data and code availability

All primary input data used in this study are publicly available. This includes the NHANES participant variables, NHANES oral microbiome datasets, alpha- and beta-diversity metrics, and the GOLD-DB prokaryotic annotations integrated for all analyses reported herein. The complete dataset needed to replicate our study has been deposited and accessible at https://doi.org/10.5281/zenodo.17849473. All code used for data processing, database integration, analyses, and generation of the tables and figures in this manuscript is publicly available at: https://github.com/terry-b-cho/OralMicroNHANES.

## Acknowledgments

We thank Christopher Smillie, Ph.D. (Assistant Professor, Harvard Medical School) for computational support and training, and Adarsh Kumbhari, Ph.D. for input on microbiome network visualization approaches. We thank Marco Jost, Ph.D. (Assistant Professor of Microbiology, Harvard Medical School) and Tami D. Lieberman, Ph.D. (Hermann L. F. von Helmholtz Career Development Professor), for insightful discussions on age-associated accumulation patterns and the interpretation of network community structure. We also acknowledge the faculty and members of the Department of Biomedical Informatics, Harvard Medical School and the Bioinformatics and Integrative Genomics (BIG) Ph.D. Program for training support. This work was supported by NIH grants NIEHS R01ES0320470, NIDDK R01DK137993, and NIEHS U24 ES036819.

## Author contributions

Conceptualization, B.C., B.T.T., C.J.P., and A.D.K.; data curation, C.J.P.; formal analysis, B.C.; statistical design, B.C. and C.J.P.; software, B.C.; visualization, B.C.; supervision, C.J.P., B.T.T. and A.D.K.; project administration, B.T.T. and C.J.P; writing – original draft, B.C. and B.T.T. and C.J.P.; writing – review and editing, B.C., B.T.T., C.J.P., and A.D.K.; C.J.P. served as the primary supervisor for the project.

## Declaration of interests

A.D.K. is an advisor at FitBiomics. B.T.T. consults for Seed Health and Enzymetrics Biosciences on microbiome study design and analysis.

## STAR★Methods

### Key Resources Table

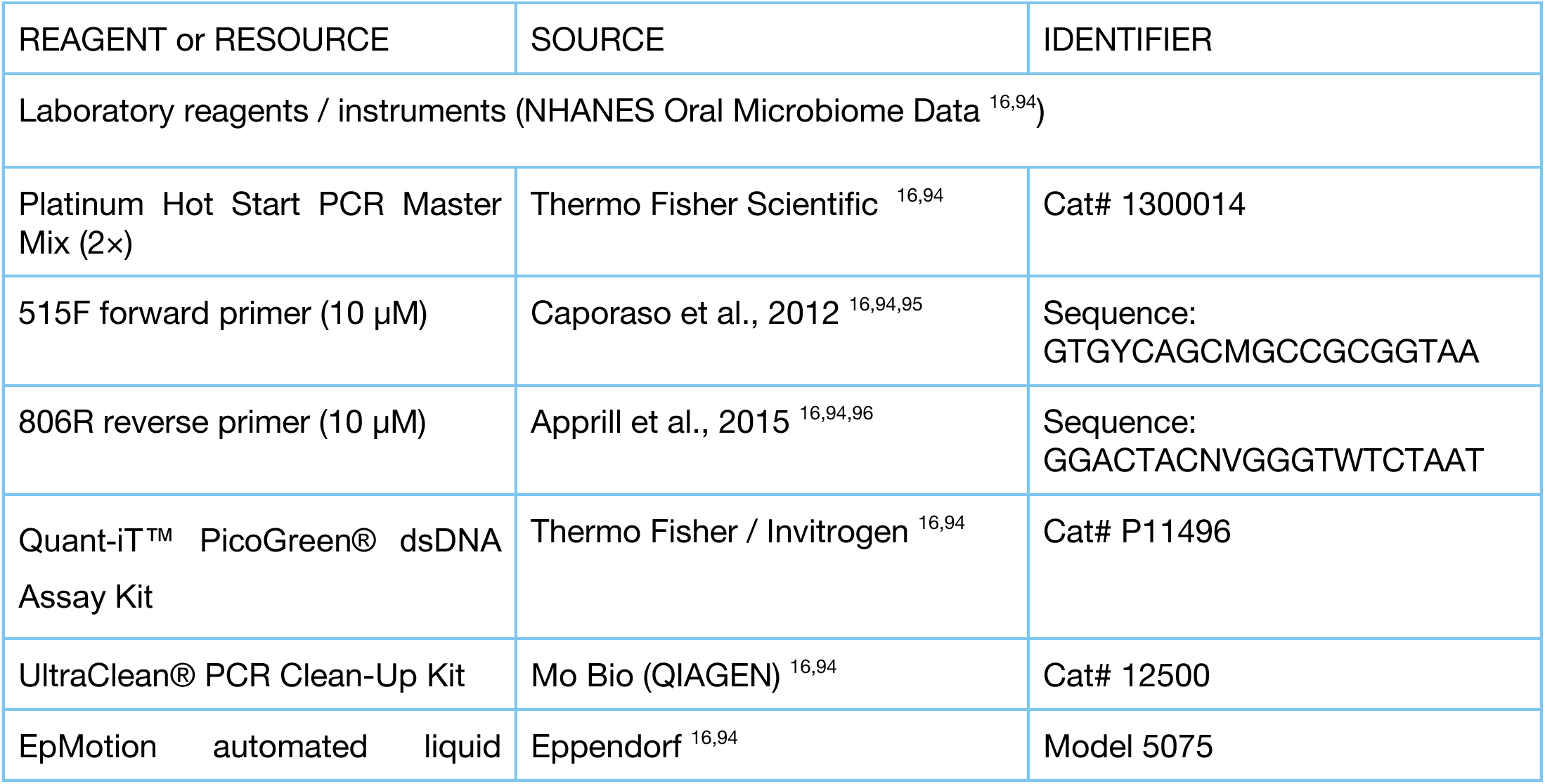
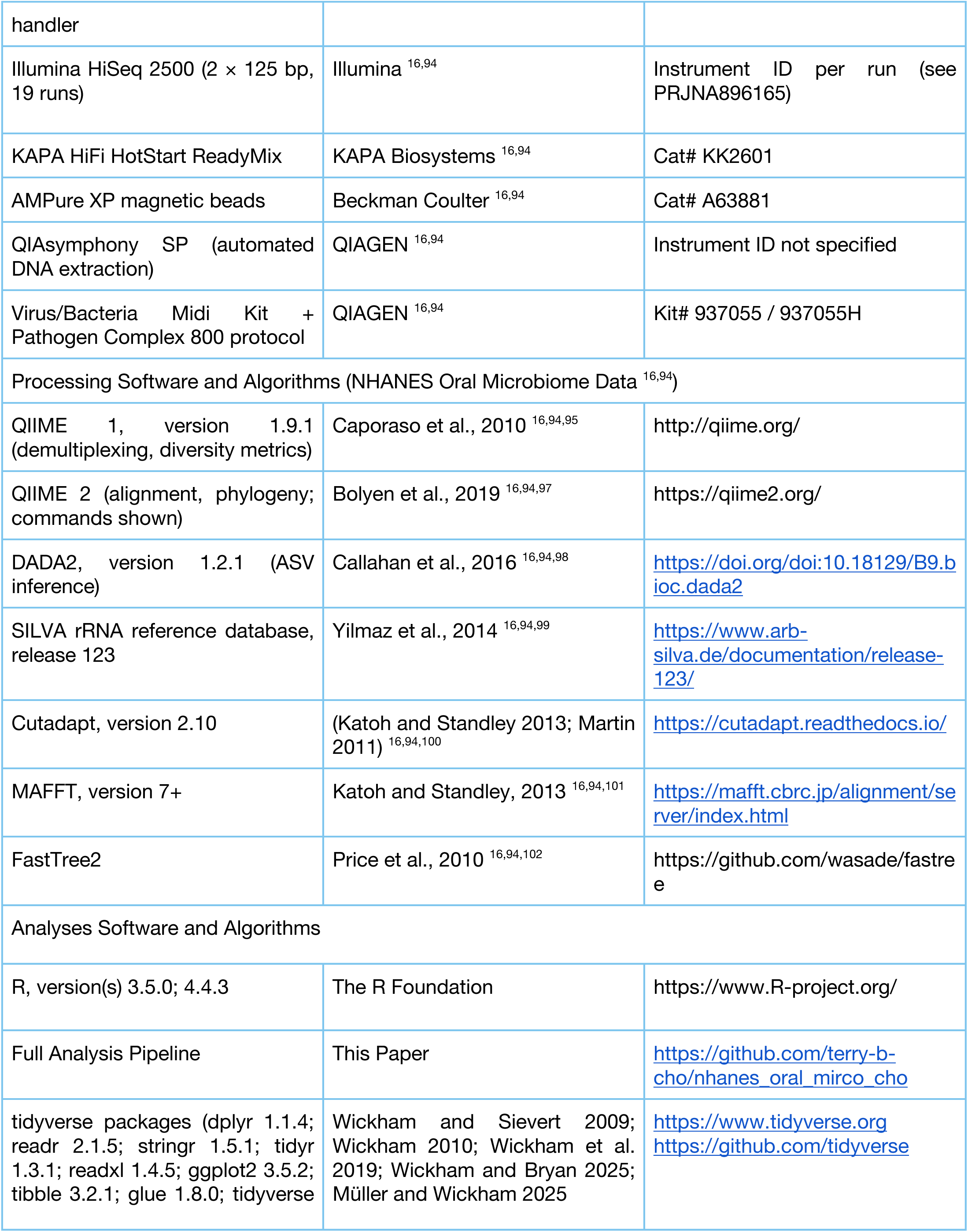
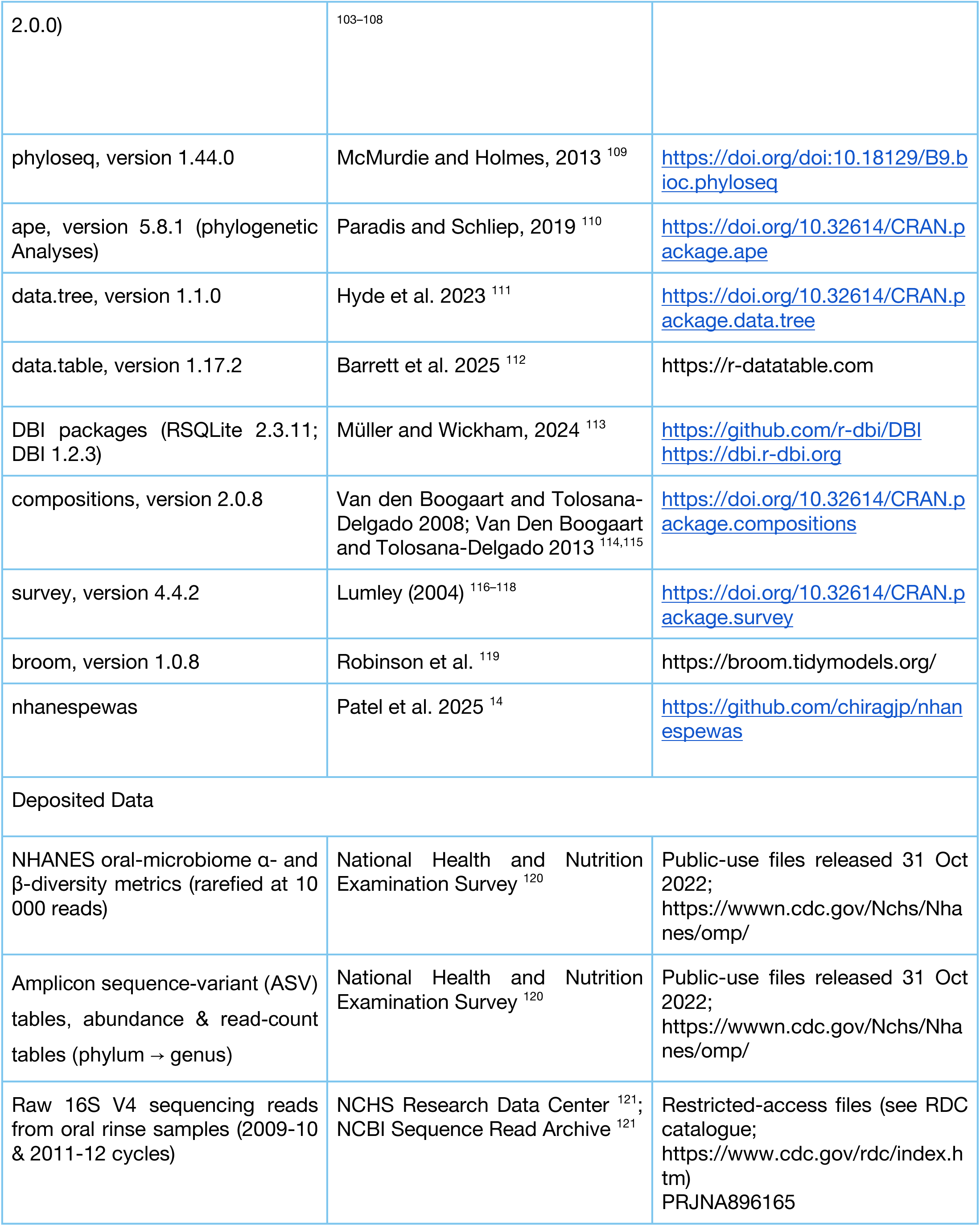
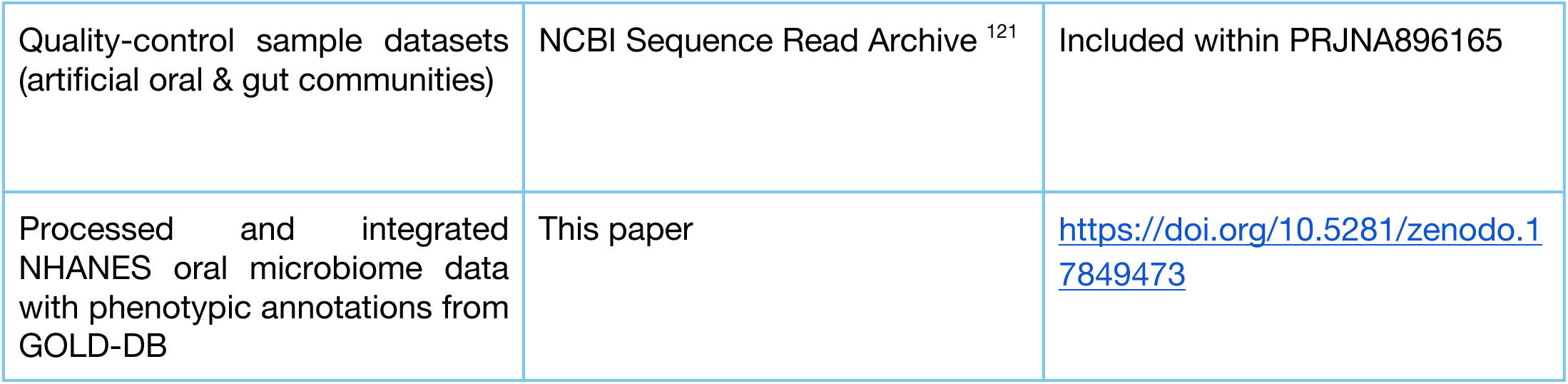

### Experimental model and study participant details

#### National Health and Nutrition Examination Surveys (NHANES) framework and oral microbiome data collection

##### NHANES 2009-2010 and 2011–2012 participant eligibility and biospecimen collection for sequencing

NHANES employs a four-stage sampling design to represent the non-institutionalized civilian population of the United States: (i) primary sampling units (PSUs), defined as counties or groups of counties, are first selected; (ii) clusters of households are then chosen within each PSU; (iii) households are randomly selected from these clusters; and (iv) one or more participants are selected from within the chosen households^122,123^. To improve estimation precision, NHANES oversampled Hispanic, non-Hispanic Black, non-Hispanic Asian, low-income non-Hispanic White/Other, and elderly (≥80 y) non-Hispanic White/Other individuals. In NHANES 2009–2010 and 2011–2012, 13,272 and 13,431 individuals were selected for screening, of whom 10,537 and 9,756 were interviewed, and 10,253 and 9,338 were examined in the Mobile Examination Center (MEC), respectively ^124^. Protocols were approved by the NCHS Ethics Review Board with written informed consent from all participants.

All examined NHANES participants aged 14 to 69 during the consecutive 2009–2010 and 2011–2012 cycles attending a mobile examination centre (MEC) were eligible for the oral rinse protocol, originally implemented to investigate the epidemiology and type distribution of human papillomavirus (HPV) infection in the U.S. population ^15^. Oral rinse specimens were collected using 10 mL of sterile saline or Scope™ mouthwash, as detailed in Chapters 10–12 of the NHANES Laboratory/Medical Technologists Procedures Manual ^125^. In 2009–2010, specimens from 4,847 participants were evaluated for HPV, while 455 were not evaluated and the remaining were not collected. In 2011–2012, 4,410 specimens were evaluated, with 386 not evaluated and the rest not collected ^126,127^. These evaluated oral rinse specimens were used to perform oral microbiome sequencing ^16^.

##### 16S rRNA Microbiome sequence processing and taxonomic assignments ^15,16,94^

DNA was extracted using the QIAsymphony SP (QIAGEN) with the Virus/Bacteria Midi Kit and Pathogen Complex 800 protocol across 72 plates (2009–2010) and 60 plates (2011–2012), each including an extraction blank, synthetic oral, and gut community controls. PCR amplification was performed in triplicate 25 µL reactions using either 2× Platinum™ Hot-Start Master Mix (Thermo Fisher) with 0.5 µM 515F/806R primers and 1 µL template DNA, or 2× KAPA HiFi HotStart ReadyMix (KAPA Biosystems) with 0.2 µM 515F/806R primers containing Illumina overhangs and 5 ng input DNA. Thermocycling followed standard protocols (Platinum: 94 °C 3 min; 35 cycles of 94 °C 60 s, 50 °C 60 s, 72 °C 105 s; KAPA: 95 °C 3 min; 25 cycles of 98 °C 20 s, 55 °C 15 s, 72 °C 30 s; both with final 72 °C extensions). Amplicons were pooled, quantified (PicoGreen®, Thermo Fisher), purified (UltraClean® PCR Clean-Up or AMPure XP beads, Beckman Coulter), indexed (8 cycles), and sequenced on the Illumina HiSeq 2500 (2 × 125 bp, rapid mode). Runs or lanes with <100 reads were repeated; only forward reads were retained due to low-quality reverse overlap.

Adapters were trimmed with Cutadapt v2.10, and reads were demultiplexed using QIIME 1 v1.9.1 (Phred ≥20). Due to insufficient overlap, only forward reads (125 bp) were processed using DADA2 v1.2.1 (parameters: maxEE=2, truncLen=120, truncQ=2, pooling=“pseudo”), yielding 41,378 ASVs across 10,442 samples. Taxonomy was assigned using the SILVA r123 naïve Bayes classifier trained on the 515F/806R region. One mitochondrial ASV (SV1032) was removed; RSV and RB filtered tables were generated. ASVs were aligned using MAFFT, and a midpoint-rooted phylogeny was constructed with FastTree2 via QIIME 2 v2023.2. Alpha diversity (observed richness, Faith’s PD, Shannon, Simpson) was computed from ten rarefied tables (2,000–10,000 reads), and β-diversity (Bray–Curtis, unweighted and weighted UniFrac) was calculated using the 10,000-read rarefied tables. Taxonomic summaries at all ranks were generated without rarefaction using QIIME 1.

All NHANES oral microbiome data used in this study were not produced as part of this manuscript. Data described here were prepared and generated by Vogtmann et al. (2023) and are publicly available; raw FASTQ files and restricted variables can be accessed via the NCHS Research Data Center upon request^16,121^.

##### Data Transformation and Statistical Strategy for Oral Microbiome-Wide Association Analysis

###### Data normalization and transformation strategy

Since survey-weighted logistic and linear regressions rely on assumptions of linear additivity in Euclidean space and approximate independence among predictors ^128^, we reasoned that simple transformations optimizing machine-learning accuracy may inflate error rates in design-weighted regression scenarios. For example, compositionally aware log-ratio transformations—mainly the centred log-ratio (CLR)—map the D-part (D−1) compositional simplex to real Euclidean space, allowing unbiased inference in linear or logistic regression and forming the backbone of log-contrast methods like LOCOM, which uses clr coordinates within a penalised logistic model to control the false-discovery rate ^129,130^.

Given complementary strengths and the absence of consensus ^131,132^ on a single optimal normalization method for inference in survey-weighted regression schema, we pre-specified four analysis scenarios— (i) untransformed relative counts, (ii) CLR to interrogate the robustness of association signals in NHANES oral microbiome data. Our study included oral microbiome data from 4,960 participants in the 2009–2010 NHANES cycle (“F”) and 4,887 from the 2011–2012 cycle (“G”), yielding a pooled sample size of 9,847 individuals. We analyzed 1,349 operational taxonomic units (OTUs), spanning taxonomic ranks from phylum to genus. As OTU vectors reside in the compositional simplex, we applied appropriate transformations prior to survey-weighted linear or logistic regression modeling to account for compositional constraints.

Raw read-count *M_i,m_* and their corresponding relative-abundance forms #_%,’_ matrices were merged with the complete NHANES tables and stored into a single SQLite archive (Can be found in: https://doi.org/10.5281/zenodo.17849473). For each of 5 taxonomic ranks (Phylum, Class, Order, Family, Genus) and for each data cycle 2009-2010 (F) and 2011–2012 (G), we generated transformed versions of these tables as independent tables.

###### Transformation definitions

All transformations were applied independently for each taxon before model fitting as followed:

For every participant *i* ∈ {*1*, ⋯, *N*)}, genus-level counts vector *M^i^* and for each genus (taxa) *m* ∈ {*1*, ⋯, *D*}:

1. Relative abundance (*P_i,m_* Proportions):

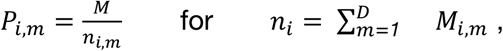
2. Centred log-ratio (*CLR*(*M_i_*_,*m*_)):

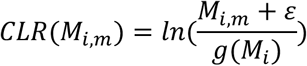

where the geometric mean 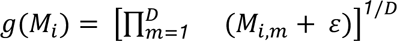 mapping the D-part compositional simplex to *R^D^*^-1^ after adding a pseudo-count ε = *1* for a sparse data and to avoid log *0*.

###### NHANES survey design specification

We incorporated the complex survey design as specified by NCHS Technical Documentation and protocols published for the years 2007-2010 ^123^ and 2011-2014 ^122^. Survey design objects were declared with stratification (SDMVSTRA; primary sampling units), clustering (SDMVPSU; masked variance strata), and survey weights (WTMEC2/4YR; MEC weights) using the *survey* v4.4-2 R package. For each variable pair (*X*, *Y*), we identified all cycles with data available for both variables (*C_Y_*_,*X*_∈ {*F*, *G*}). If both cycles were available, we performed pooled-cycle analysis; if only one cycle was available, we used single-cycle analysis. For pooled analyses, we constructed NCHS-compliant 4-year weights (WTMEC4YR = WTMEC2YR/2) and generated unique design identifiers to avoid overlap across cycles.

###### Survey-weighted association modelling frameworks

As specified in Table 1, five analysis types were defined to characterize microbiome–host associations across distinct microbial genus to host variable pairings at the genus level. The *CLR*(*M_i_*_,*m*_) values entered. Covariate set *C_i_* comprised of fully expanded 13 variables of age, gender, race/ethnicity, education, nativity (US born) and poverty-to-income ratio as specified in Figure 1.

For demographic variables or exposure variables *X_i_*, clr-transformed abundances were modelled as outcomes:

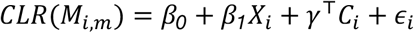

For measurable host phenotype variables *Y_i_*, *clr* abundances were modeled as predictors:

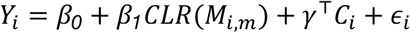

Oral conditions and host disease outcome *Y_i_*, were defined as binary variables and were modelled using a logit link:

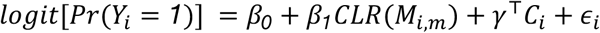

These models incorporated NHANES-specific strata, clusters, and sampling weights (as defined in the NHANES survey design specification), ensuring that microbiome normalization procedures remained orthogonal to the survey design. For each host-variable-microbiome pair (*h*, *m*), we estimated the survey-weighted effect sizes (regression coefficients), robust standard errors 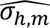 based on Taylor linearization, associated p-values, and survey-weighted R² values. Multiple-testing correction was performed using both the Benjamini–Hochberg^133^ false discovery rate method and Storey’s^134^ q-value approach. All regression outputs are deposited and available at https://doi.org/10.5281/zenodo.17871009.

###### GOLD DB Annotation integration

To enrich genus-level taxonomic data from the NHANES oral microbiome with functional context, we integrated microbial phenotypic annotations from the Joint Genome Institute’s Genomes OnLine Database^135^ (GOLD; v.10), which catalogs phenotypic and ecological metadata for more than 400,000 microbial isolates and metagenome-assembled genomes. Species-level traits available in GOLD – including oxygen requirement, Gram-stain reaction, motility, sporulation, and selected metabolic characteristics– were aggregated to the genus level to enable direct integration with NHANES genus-level 16S rRNA profiles (SILVA v123). This mapping produced a functional annotation framework encompassing 1,349 NHANES OTUs corresponding to 4,215 GOLD-annotated genera.

Genus-level harmonization was required to maximize cross-database mapping accuracy. NHANES genus names were standardized using a parsing algorithm that removed sub-group designations, and other non-canonical modifiers, followed by case normalization and synonym resolution. Canonicalized names were then mapped to GOLD entries via exact string matching to preserve taxonomic specificity; unmapped genera were retained as missing values for downstream analysis. Across all classifiable NHANES oral genera (i.e., excluding “unclassified” taxa), 85.06 % were successfully matched to at least one GOLD-annotated microbial entry. Microbial trait/phenotype variables were encoded quantitatively. Binary traits (Gram-positive, motility, sporulation) were assigned values of 0 (absent; gram-negative; non-sporulating) or 1 (present; gram-positive; sporulating), and genus-level indices were computed as the mean of all constituent species (range 0-1). Oxygen requirement was recorded as a continuous ordinal scale (anaerobe = 0, microaerophilic = 0.25, facultative = 0.5, aerobe = 1.0) and averaged within genera to yield an “oxygen index.” Categorical traits were summarized as frequency distributions across species. Agreement among species within a genus was classified as high (100 % concordant), medium (≥70 % sharing the dominant trait), or low (<70 %), providing a confidence score for trait reliability. Full resulting annotations and mapping summaries are available in Table S1 (Table_S1_GOLD_Mapping_and_Index_aggregation.xlsx).

###### Compositional Differential Abundances

Compositional differential-abundance heatmaps were generated to provide an intuitive, non-regression-based comparison of group differences, offering a direct complement to the survey-weighted regression results. Labelled genera present in at least 1% of specimens or non-zero variance in CLR were retained for analysis. For group comparisons, two-sided Wilcoxon rank-sum tests were applied to binary variables and Kruskal–Wallis tests to variables with more than two levels; P values were adjusted using the Benjamini–Hochberg procedure (FDR ≤ 0.05). Significant taxa were ranked by the variability of group means, and the top 30 (or fewer, if fewer passed the significance threshold) were selected for visualization. Each heatmap panel displays the difference from the reference group using modality-specific scales: relative-abundance z-scores for raw data (capped at –1.5 to +1.5), log₂ fold change for CLR values (capped at –2.5 to +2.5). Taxa were hierarchically clustered using Euclidean distance.

###### Microbial Signature Correlation

For each survey design model fit, yielding an estimate 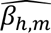 and standard errors 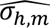, we retained taxa observed in at least 1% of the population then filtered for case imbalance for binary host variables, then filtered cases where (*FDR* ≥ *0.05*) and (*q* ≥ *0.05*). For each host variable *h*, the remaining coefficients were collected into a microbial signature vector 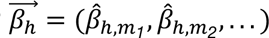 and its matching standard error vector 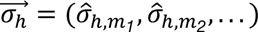, together representing the microbial association profile for that host variable *h*. Pairwise similarity between host signatures were quantified using inverse-variance-weighted host-variable correlations. For every two host variables *h_i_* and *h_j_* sharing at least three tax we computed the weight *w_m_* and correlation *r_i,j_* as followed:

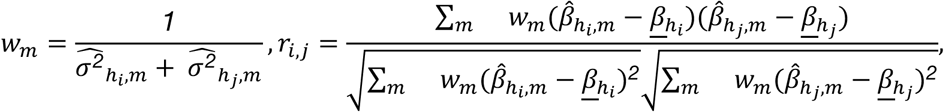

Where, 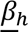 is the weighted mean of 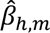. We computed a t-statistic and its p-values from:

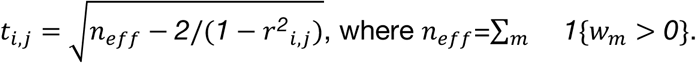

We adjusted our p-values via the Benjamini-Hochberg procedure to yield matrices of microbial signature correlations *R* = [*r_i_*_,*j*_] and *FDR* = [*FDR_i,j_*].

###### Microbiome Network Plotting

For each host variable *h* and a set of CLR transformed microbiome variables *M*, we applied an inclusion criteria of 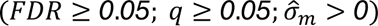 and prevalence >10%. We also applied a case-balance filter requiring cases to comprise at least 0.5% of the total sample. Let *M*^∗^ denote the resulting subset of retaining associations. We define as *N_h_* be the set of microbial nodes that appear in at least one retained association with host variable *h*. For signed regression coefficient β*_m,h_* for *M*^∗^ with respect to host variable *h*, the edge weight between all unordered pairs of distinct microbes (*m_i_*, *m_j_*) is defined as 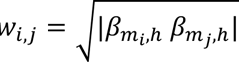 reflecting the joint magnitude of two independent host associations. An edge is included in *E_h_* if and only if *w_i,j_* > *0*.

Using this set, we construct an undirected weighted graph *G_h_* = (*N_h_*, *E_h_*), where edges encode similarities of statistically significant host-microbe associations rather than biological interaction between microbes. Networks are only constructed when at least two retained associations and at least two distinct microbial taxa are present. To encode agreement in effect direction, we define a directional concordance indicator *c_i,j_* as *c_i,j_* = *1* (concordant positive) for 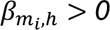 and 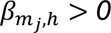 (concordant negative) if 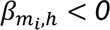 and 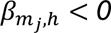, *c_i,j_* = *0* and *c_i,j_* = *0*.*5* otherwise (discordant). This quantity is used solely for visualization and does not affect graph topology.

###### Age-resolved oral microbiome clustering with data-driven cluster determination

To characterize age-related patterns in oral microbial community structure, we analyzed genus-level relative abundance profiles from 9,349 NHANES participants spanning 11 age groups (14–19, 20–24, 25–29, 30–34, 35–39, 40–44, 45–49, 50–54, 55–59, 60–64, and 65–69 years). Genera detected in at least 10 % of participants (93/965) were retained to ensure robust prevalence. For each genus, we computed the mean relative abundance within each age group and standardized values (z-scores) across the 11 groups, producing a 93 × 11 matrix representing genus-specific abundance trajectories through age.

We applied hierarchical clustering (Ward’s minimum-variance linkage on Euclidean distances) to identify genera with similar age trajectories. To determine the optimal number of clusters, we evaluated candidate k values (2–9) using two complementary methods: (i) silhouette maximization, selecting k that maximized mean silhouette width as a direct measure of within-cluster cohesion versus between-cluster separation; and (ii) the gap statistic, comparing observed within-cluster dispersion with 100 bootstrap null reference distributions generated from uniformly sampled data in 10-component PCA space. The silhouette-width diagnostic curve (Fig. 2C) showed monotonic improvement from *k = 2* (1.086) to *k = 9* (2.851). Given the higher internal consistency and interpretability of the silhouette criterion, k = 9 was adopted as the final cluster solution.

We evaluated differential abundance across age groups using Kruskal–Wallis tests for each genus, with p-values adjusted by the Benjamini–Hochberg procedure (FDR < 0.05). Visualizations included parallel-coordinate plots of all genera and per-cluster trajectories, silhouette and gap diagnostic plots, PCA and t-SNE embeddings (Rtsne; perplexity = 30, seed = 42), and a heatmap of significant genera (n = 77) with hierarchical row clustering and phylum- and cluster-level annotations. Cluster validity was confirmed by distance-to-centroid analysis and PERMANOVA (9,999 permutations) on the standardized matrix, demonstrating significant non-random grouping. The t-SNE projection (variance explained: 53.35 % and 46.65 % in dimensions 1 and 2, respectively) showed well-defined cluster neighborhoods consistent with the hierarchical solution. Cluster assignments and intermediate outputs are provided in (Supplementary Tables S8 and S9).

## Notes

https://doi.org/10.5281/zenodo.17849473

### Competing Interest Statement

The authors have declared no competing interest.

